# Defining and Evaluating Microbial Contributions to Metabolite Variation in Microbiome-Metabolome Association Studies

**DOI:** 10.1101/402040

**Authors:** Cecilia Noecker, Hsuan-Chao Chiu, Colin P. McNally, Elhanan Borenstein

## Abstract

Correlation-based analysis of paired microbiome-metabolome datasets is becoming a widespread research approach, aiming to comprehensively identify microbial drivers of metabolic variation. To date, however, the limitations of this approach have not been comprehensively evaluated. To address this challenge, we introduce a mathematical framework to quantify the contribution of each taxon to metabolite variation based on uptake and secretion fluxes. We additionally use a multi-species metabolic model to simulate simplified gut communities, generating idealized microbiome-metabolome datasets. We then compare observed taxon-metabolite correlations in these datasets to calculated ground-truth taxonomic contribution values. We find that in simulations of both a model 10-species community and of complex human gut microbiota, correlation-based analysis poorly identifies key contributors, with extremely low predictive value despite the idealized setting. We further demonstrate that the predictive value of correlation analysis is strongly influenced by both metabolite and taxon properties, as well as exogenous environmental variation. We finally discuss the practical implications of our findings for interpreting microbiome-metabolome studies.

**Importance:** Identifying the key microbial taxa responsible for metabolic differences between microbiomes is an important step towards understanding and manipulating microbiome metabolism. To achieve this goal, researchers commonly conduct microbiome-metabolome association studies, comprehensively measuring both the composition of species and the concentration of metabolites across a set of microbial community samples, and then testing for correlations between microbes and metabolites. Here, we evaluated the utility of this general approach by first developing a rigorous mathematical definition of the contribution of each microbial taxon to metabolite variation, and then examining these contributions in simulated datasets of microbial community metabolism. We found that standard correlation-based analysis of our simulated microbiome-metabolome datasets identifies true contributions with very low predictive value, and that its performance depends strongly on specific properties of both metabolites and microbes, as well as on the surrounding environment. Combined, our findings can guide future interpretation and validation of microbiome-metabolome studies.

## Introduction

Microbial communities have a tremendous impact on their surroundings, ranging from the degradation of environmental toxins (1) to the production of climate change-relevant metabolites (2). Host-associated communities, in particular, have a substantial impact on their hosts, and often produce a diverse set of metabolites that interact with numerous host pathways. In humans, such microbiome-derived metabolites have been identified as contributing factors to a wide array of diseases including heart disease (3), autism (4), non-alcoholic fatty liver disease (5), colon cancer (6), inflammatory bowel disease (7), and susceptibility to infection (8). Characterizing the ways microbial communities modulate their environments and the relationship between community structure and metabolic impact is therefore a major, timely, and complex challenge with promising implications for human health, as well as to environmental stewardship, agriculture, and industry.

When facing this challenge, perhaps the most important task is identifying specific community members that drive variation in metabolites of interest. Taxa responsible for observed metabolic differences across communities may be ideal targets for interventions aiming to modify metabolic phenotypes. Their identification, however, can be a daunting task. Complex microbial communities are often composed of hundreds or thousands of poorly characterized species, each with a unique and frequently unknown complement of metabolic capacities. Even when multiple species are known to possess the potential to synthesize or degrade a metabolite of interest, the metabolic activity of each species (and consequently, its contribution to metabolic variation) may be different (9). Moreover, community ecology, interspecies interactions, and nutrient availability (e.g., via diet) can all regulate and influence the metabolic activity of each species, rendering the link between community members and metabolic products extremely complex and challenging to infer (10–12).

To address this challenge and to identify community members that play an important role in metabolic variation, a growing number of studies are now comprehensively assaying multiple facets of community structure across samples, including, most notably, taxonomic and metabolite compositions (13). For example, to investigate the links between taxonomic shifts and metabolic phenotypes in the healthy vaginal microbiome and in bacterial vaginosis, a recent study used a combination of 16S rRNA qPCR, sequencing, and both global and targeted metabolomics (14). Another study, aiming to identify taxonomic and metabolic features of resistance and susceptibility to *C. dificile* infection in the mouse gut similarly applied 16S rRNA sequencing and global metabolomics (15). In another example, researchers characterized metabolic and microbial features of periodontitis in the oral microbiome before and after treatment, combining 16S rRNA sequencing, shotgun metagenomic sequencing, and metabolomics (16). These are just a few examples of a plethora of recent microbiome-metabolome studies, investigating the metabolic effects of microbiome variation in the contexts of chronic and infectious disease, agriculture, precision medicine, nutrition, fermented food science, and more (17–24). Such multi-omic studies are also a major focus of several large-scale initiatives to study both host-associated and environmental microbiomes (25, 26).

Given the taxonomic and metabolomic profiles obtained via such microbiome-metabolome assays, the vast majority of studies rely on simple univariate correlation-based analyses to link variation in community ecology to variation in metabolic activity (11, 14, 15, 27–30). Such analyses specifically aim to identify species whose abundance across samples is correlated with the concentration of metabolites, often assuming that highly significant correlations reflect a direct mechanistic link between the taxon and metabolite in question. These studies further regularly assume that positive correlations imply synthesis and negative correlations imply degradation, or that targeting the microbe in question could be used to modulate the concentrations of the metabolites with which it is correlated. For example, a recent study characterizing the microbiome and metabolome in Spleen-yang-deficiency syndrome (29) concluded that a positive correlation between Bacteroides and mannose likely resulted from extracellular degradation of mannan into mannose by that taxon. Similarly, a study of antibiotic perturbations to the microbiome and metabolome stated that the presence of several weak positive and negative correlations between genera and arginine supported the conclusion that arginine levels may be affected by many community members with high functional redundancy (27).

Yet, to date, the extent to which a correlation-based analysis effectively detects direct metabolic relationships between taxa and metabolites is unclear. Obviously, a strong correlation between the abundance of a certain species and the concentration of a metabolite across samples *could* reflect direct synthesis or degradation of the metabolite by that species, but could also arise due to environmental effects, precursor availability, selection, random chance, or co-occurrence between species. Similarly, cross-feeding, external host processes, and varying enzymatic regulation can mask a correlation even when this species does in fact contribute to observed metabolite variation. Indeed, previous studies have suggested that microbe-metabolite correlations must have a high rate of false positives (31), and a recent experimental study pairing microbiome-metabolome correlation analysis with *in vitro* monoculture validations found anecdotally that several observed correlations were in fact false positives (32). The limitations of correlation analysis have also been discussed and well-characterized in other data types (for example (33, 34)). Importantly, however, the extent of such limitations in the context of microbiome-metabolome studies, the way they are shaped by microbial community metabolism, and their impact on data interpretation in this context have not been systematically evaluated.

Importantly, two crucial challenges hinder a comprehensive and systematic evaluation of correlation-based analysis. The first is the lack of a rigorous general definition of a microbe’s contribution to metabolite variability. While establishing the main taxonomic contributors to metabolite variation may be straightforward for specialized, well-characterized metabolites that are synthesized by just a single taxon, it can be much less clear for metabolites that can be synthesized (and/or degraded or modified) by many different taxa in the community. The second challenge is the absence of ground truth data on the nature of microbe-metabolite relationships. While limited data on the taxa driving metabolite shifts can be obtained from comparative mono- and co-culture studies (32, 35, 36), large-scale and comprehensive datasets that link species and metabolite abundances in the context of a complex community, for which the precise impact of each species on observed metabolite variation is known, are currently not available.

In this study, we address these two challenges, combining a novel framework for quantifying microbial contributions with a model-based simulated dataset. Specifically, we first introduce a generalizable and rigorous mathematical framework for decomposing observed metabolite variation and quantifying the contribution of each community member to this variation based on uptake and secretion fluxes. Second, we use a dynamic multi-species genome-scale metabolic model to simulate the metabolism of microbial communities of varying complexity and to generate idealized datasets of paired taxonomic and metabolomic abundances, with complete information on metabolite fluxes, microbial growth, interspecies interactions, and environmental influences. Applying our mathematical framework to these simulated datasets, we could then compare calculated contribution values to observed taxon-metabolite correlations and evaluate the ability of correlation-based analyses to identify key microbial contributors. We were additionally able to investigate factors that shape the relationship between community composition and metabolism in depth and to identify specific properties and mechanisms that impact the performance of microbiome-metabolome correlation studies.

Notably, given the objectives of this study, we intentionally focus on characterizing microbiome-metabolome relationship in a model-based, tractable, and well-defined setting. Indeed, our metabolic model may not perfectly capture all the complex and diverse mechanisms that are at play in host-associated communities; however, considering the scope of this study, accurately modeling the metabolism of a specific community may not be crucial. Rather, for our analysis, we want our simulated data to recapitulate broad trends observed in naturally occurring microbial ecosystems, as indeed has been observed in similar models (37–41). Moreover, utilizing this model-based approach allows us to dissect the relationship between community composition and metabolic phenotypes without the complexities inherent to *in vivo* communities (including spatial heterogeneity, measurement error, inter-microbial signaling, or strain-level variation), and with variation in the concentrations of environmental metabolites resulting exclusively from microbial metabolic activity. Analyzing the ability of a correlation-based analysis to detect true microbial drivers of metabolite variation in these simplified, best-case settings provides a baseline for the expected performances of such analyses in real microbiome-metabolome studies.

## Results

### Quantifying the impact of individual microbial species on variation in metabolite concentrations

In this study, we consider a microbial community as an idealized system, consisting of a population of multiple microbial species in a shared, well-mixed, biochemical environment. Each species uptakes necessary metabolites from the shared environment, performs a variety of metabolic processes to promote its growth, and secretes certain metabolites back into the shared environment. We additionally assume that certain nutrients flow into the environment and that microbial cells and metabolites are diluted over time. These processes can represent, for example, the inflow of dietary nutrients and the transit through the gut in the context of the gut microbiome. For simplicity, we primarily consider a constant inflow and dilution rate, as in a chemostat setting. Accordingly, a microbiome-metabolome study can be conceived as analyzing a set of several such communities (at a certain point in time), each with a different composition of microbial species and correspondingly variable environmental metabolite concentrations. We focus initially on a controlled setting with identical nutrient inflow across all microbiomes, but later examine the impacts of differences in nutrient inflow between communities.

Given this setting, we first sought to establish a rigorous and quantitative framework for defining the impact of each microbial species (or any taxonomic grouping) in the community on the variation observed in the concentration of a given metabolite across community samples. We focused on species that *directly* modulate the environmental concentration of a given metabolite via synthesis or degradation, ignoring indirect effects via, for example, the synthesis of a precursor substrate that could impact the metabolic activity of other species. We noted that the total concentration of a metabolite in the environment can be represented as the sum of cumulative synthesis or degradation fluxes of this metabolite by each of the *n* species in the community, as well as cumulative environmental fluxes (e.g., total nutrient inflow and dilution). Formally, the metabolite concentration, *M*, can therefore be expressed as a sum of *n* dependent random variables *m_i_*, where each *m_i_* denotes the overall synthesis or degradation of the metabolite by each species, along with an additional random variable *m_env_*, denoting the overall impact of environmental processes.

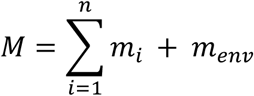

As discussed above, when analyzing microbiome-metabolome datasets, the goal is often to identify taxa responsible for *changes* in the concentration of a metabolite of interest across a set of samples. Accordingly, here we wish to quantify the *contribution* of each species to the *variance* in the concentration of that metabolite across samples. Specifically, in the formulation above, *var*(*M*) depends on the variance in the constituent microbial and environmental factors, as well as the covariance between these components. This variance can then be linearly separated into *n*+*1* terms, representing the contribution of each species (denoted *c_i_*), and of any environmental nutrient fluxes (denoted *c*_*env*_) to the total variation in the metabolite:

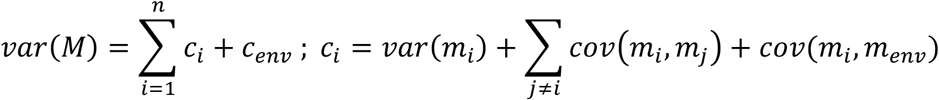

If the nutrient inflow is constant across samples, its effect can be ignored and its contribution to the variance *c_env_* is 0. Additionally, in a chemostat setting, the dilution of each metabolite can be accounted for in the calculation of each contribution, as it depends strictly on the dilution rate and on previous metabolite concentrations (Methods). Finally, in order to compare species contributions across metabolites and to represents the relative share of the total variance of a given metabolite that is attributable to species *I*, we defined the *relative* contribution to variance *ĉ_i_* of each species *i* to metabolite *M* by normalizing contribution values by the metabolite’s total variance:

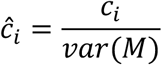

This framework for calculating microbial contribution values provides a systematic measure of the causal impact of each taxon on observed variation in the environmental concentration of each metabolite, distilling the effect of complex ecological and metabolic interactions to a concise and interpretable set of quantities. Moreover, the obtained contribution profile is a linear decomposition of observed metabolic variation, wherein the sum of contributions of all species equals the observed variation in the metabolite. Notably, when a species’ activity has large negative covariances with the activities of other community members, contribution values can be negative. Such negative contribution values indicate that a species’ secretion or uptake of that metabolite varies in a way that mitigates the activity of others. Correspondingly, contribution values can be greater than 1, reflecting scenarios in which a species in fact generates more variation of this metabolite than is ultimately observed, but that its impact is mitigated by other species.

It is also worth noting that our analytical decomposition of contributions to variance is mathematically equivalent to calculating the Shapley values for the variance in metabolite concentrations (see Methods and Figure S1). Shapley value analysis is a game theory technique that defines an individual’s contribution to a collective outcome, and has been shown to be the only general definition that is efficient, linear, symmetric, and assigns zero values to null contributors (42). A similar, Shapley value-based approach was recently applied to address the related problem of identifying the primary taxonomic contributors to differential functional abundances in metagenomic data (43).

### A multi-species metabolic model for generating complex microbiome-metabolome data

We next set out to generate a large-scale dataset of microbiome-metabolome profiles with complete information about metabolite uptake and secretion fluxes. To this end, we used a multi-species metabolic model to simulate the growth, dynamics, metabolism, and environment of a simple microbial community. This model is based on a previously introduced genome-scale framework for modeling the metabolism of multi-species communities and for tracking the metabolic activity of each community member over time (44, 45). Briefly, this framework assumes that each species optimizes its growth selfishly given available nutrients in the shared environment and predicts the metabolic activity for each species in short time increments using Flux Balance Analysis (46). After each increment, the model uses the predicted metabolic activities of the various species to update the biomass of each species and the concentration of metabolites in the shared environment (hence, potentially impacting the growth and metabolism of other species in subsequent time steps). Importantly, this model allows for the natural emergence of metabolic competition and exchange between species, as well as selection for taxa with the most efficient growth rate in a given nutrient environment. Full details of this model and simulation parameters can be found in the Methods.

We specifically modeled a simplified gut community that was previously explored experimentally (47). This community includes 10 representative gut species, spanning the major clades found in the human gut and collectively encoding the key metabolic processes taking place in this environment, including breakdown of complex dietary polysaccharides, amino acid fermentation, and removal of fermentation end products via sulfate reduction and acetogenesis. Genome-scale metabolic models of these 10 species were obtained from the AGORA collection (40) – a recently introduced set of high-quality gut-specific metabolic models. To mimic the experimental gnotobiotic mouse setting (47), we simulate growth in a chemostat, with a nutrient inflow mimicking the content of a standard corn-based mouse chow, and a dilution rate consistent with mouse transit time and gut volume. While maintaining this nutritional environment, we systematically explored the landscape of possible community compositions, varying the initial relative abundance of each species from 10% to 60% (with a consistent total abundance equal to the community carrying capacity), resulting in a total of 61 different community compositions. For the analysis below, we simulated growth for 144 hours (as 576 15-minute time steps). For most community compositions considered, this simulation time consisted of an initial stabilization period followed by a transition to a near-steady-state equilibrium with little change in community composition (Figure 1A). Notably, across the various simulations, some species maintained high abundances throughout the course of the simulation, while others reverted to lower levels.

**Figure 1.**
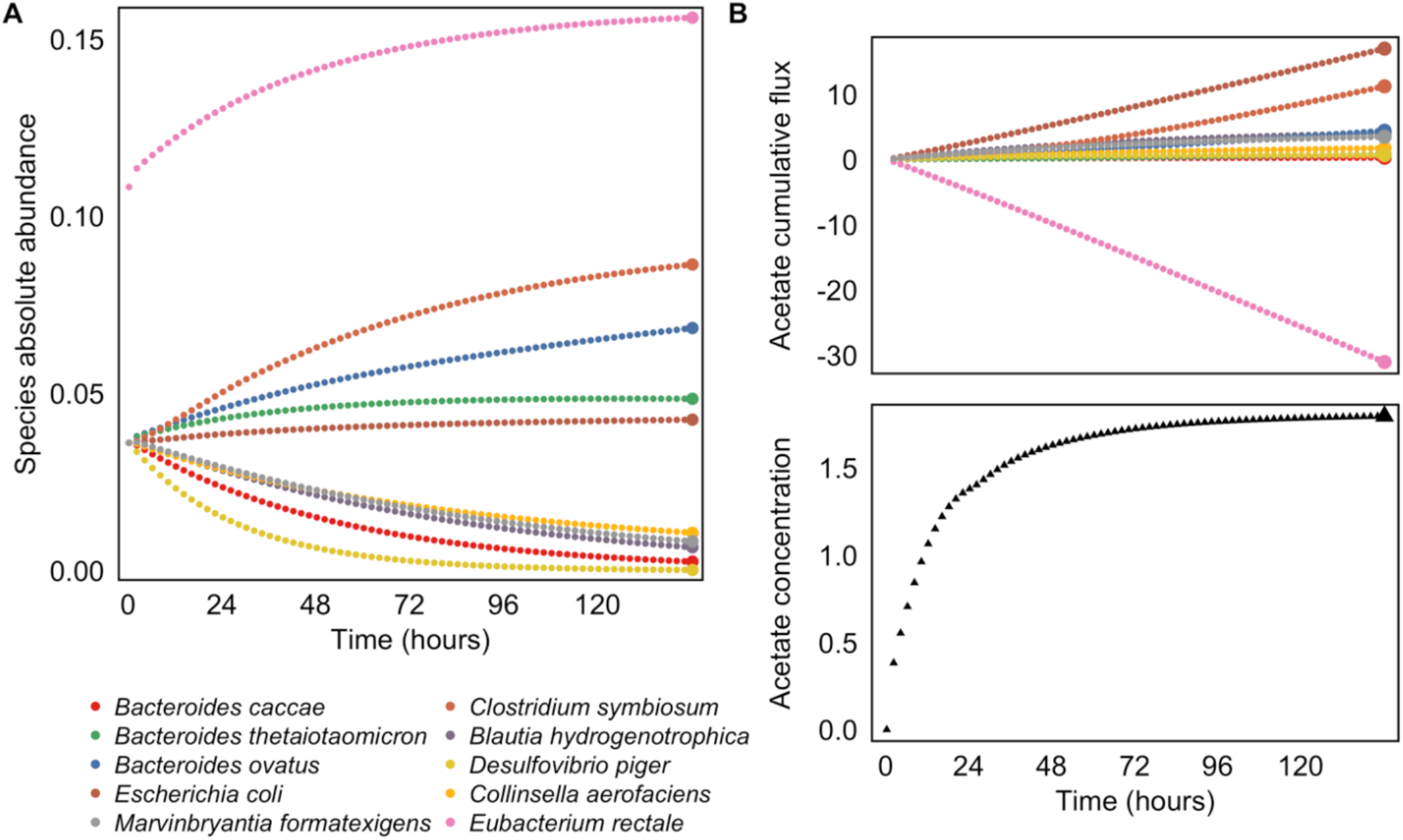
Simulating multi-omic data with a dynamic multi-species genome-scale framework. **(A)** Community species abundances throughout a single simulation run. Abundances were quantified in units of microbial biomass. In this simulation, community composition was initialized with a high relative abundance of *Eubacterium rectale*. For visual clarity, only every eighth time step is illustrated. Species abundances at the final time point (highlighted with larger colored circles) were used for calculating species-metabolite correlations. **(B)** Cumulative secretion and uptake of acetate by each community member, throughout the same simulation run illustrated in panel A. Acetate was synthesized by several species and consumed by *E. rectale* over the course of the simulation. Total cumulative fluxes (highlighted with larger colored circles) were used for calculating species contributions to metabolite variation. The bottom plot illustrates the resulting environmental concentration of acetate at each time point. The metabolite concentration at the final time point (highlighted with a larger black triangle) was used for calculating species-metabolite correlations.

Throughout the course of each simulation, we recorded the abundances of each species, the secretion and uptake rate of each metabolite by each species (as well as internal reaction fluxes), and the concentration of each metabolite in the environment (Figure 1A-B), thereby obtaining a comprehensive dataset describing species composition, metabolic activities, and metabolite concentrations across 61 different communities. To mirror the typical structure of a microbiome-metabolome cross-sectional dataset, we specifically considered the abundances of species and the concentrations of metabolites in the environment at the end of each simulation (i.e., after the final time point; see Figure 1). 60 of the 68 metabolites present in the nutrient inflow exhibited at least some variation across communities, as did 18 additional microbially-produced metabolites. Metabolite variation was generally low (median coefficient of variation 0.021), reflecting a relatively stable nutrient environment, yet 25 metabolites (32%) did have a coefficient of variation greater than 0.1. For downstream analysis, we excluded metabolites without substantial measurable variance across samples, filtering those with variance at or below the 25^th^ percentile. This resulted in a dataset of 52 variable metabolites, of which 14 are purely microbially-produced metabolites, 9 are microbially-produced but also present in the nutrient inflow, and 29 are introduced only through the nutrient inflow. Of these 52 variable metabolites, 47 are utilized by any member of the community (including 18 that are cross-fed in at least one simulation). The final species compositions and the final concentrations of several key metabolites across all simulations are shown in Figure 2A-F, and ordination plots of species and metabolite data are shown in Figure S2.

**Figure 2.**
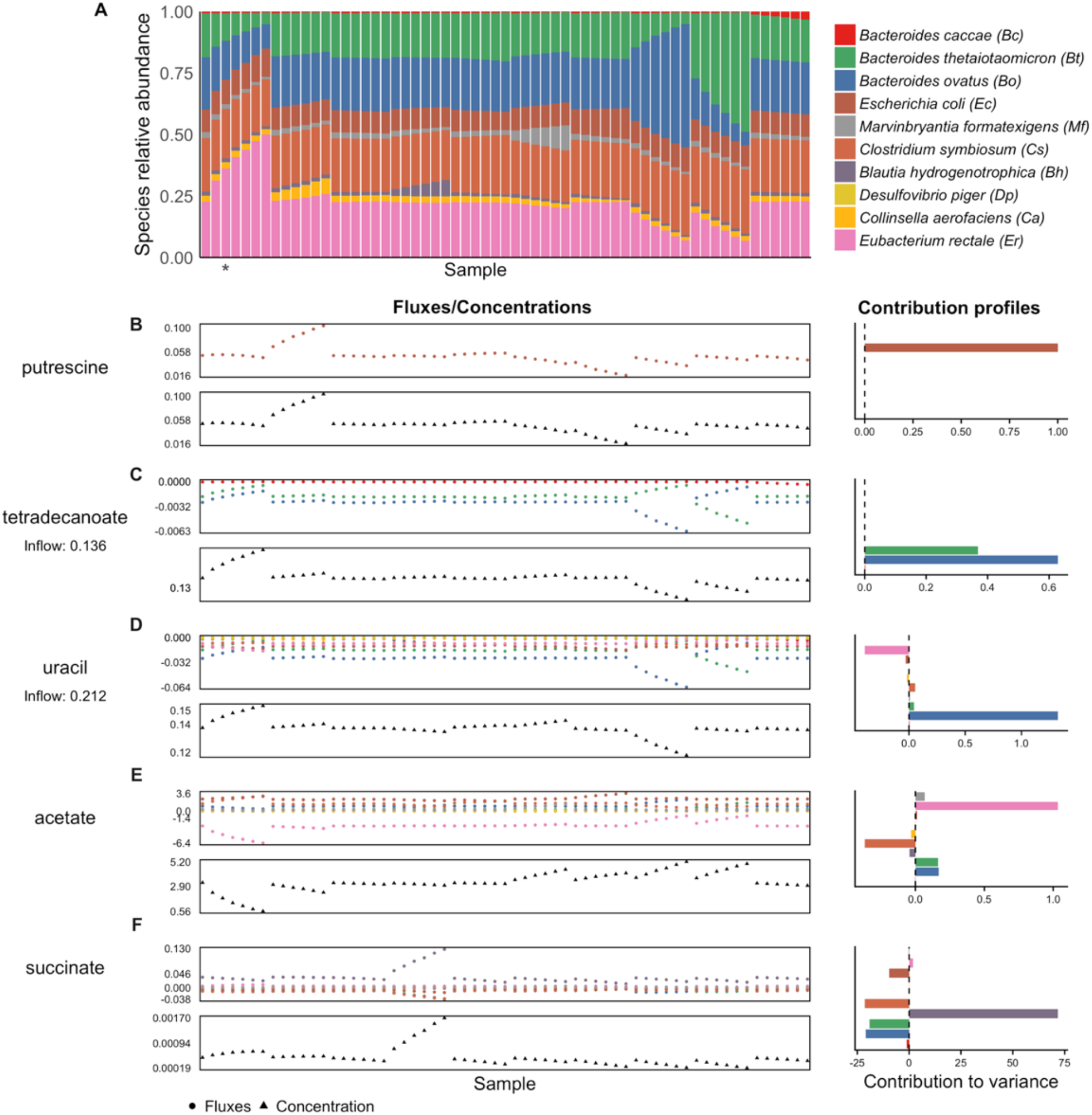
Species abundances, cumulative fluxes, and contributions to variance in metabolite concentrations in our simulated dataset. **(A)** The dataset of species abundances at the final time point of 61 simulation runs. Each bar represents a simulation run, with the colors indicating relative abundance of each species. The abundance profile from the simulation runs highlighted in Figure 1 is indicated with an asterisk. **(B-F)** For five example metabolites, the upper plot shows the total cumulative secretion or uptake of that metabolite by each species across all 61 simulation runs (or samples). The lower plot shows the corresponding environmental concentration at the final time point. The bar plot on the right shows the contribution values for each species and metabolite, calculated from the flux values and describing each species’ linear contribution to the overall metabolite variance.

Exploring this dataset, we found that species composition and metabolite concentrations exhibited complex patterns and biologically reasonable distributions (Figure S3) (49). Several metabolic processes known to occur in the mammalian gut were replicated by our simulations, including, for example, conversion of acetate to butyrate by *E. rectale* (48), and production of key microbial metabolites such as 4-aminobutyric acid (GABA), indole, and succinate. Cross-feeding relationships were observed frequently (18 metabolites), including cross-feeding of 6 amino acids, whose exchange is widespread in host-associated microbiota (50). Additionally, we ran several sets of simulations with introduced fluctuations in the nutrient inflow concentrations, and found that the resulting species compositions partially recapitulated the diet responses observed by Faith *et al.* (47) (Supplementary Results).

Clearly, the model and simulations described above represent a gross simplification of the microbiome’s structure, dynamics, and function. Importantly, however, this simplification is also an important strength. Specifically, the data obtained from these simulations provide a unique opportunity to examine the relationship between community dynamics and metabolic activity in a realistic, yet tractable model of community metabolism where complete information about the activity and fluxes of each microbial species is available (Figure S4). Indeed, our multi-species model captures many of the intricacies of bacterial genome-scale metabolism and the interconnectedness (both within and between species) of multiple metabolic processes, yet without additional complexities inherent to *in vivo* communities. Furthermore, in our simulations, variation in the concentrations of environmental metabolites results exclusively from microbial metabolic activity, with no variation in nutrient inflow or other non-microbial sources, providing a controlled setting for evaluating the relationship between community members and metabolite concentrations.

### Metabolite variation is driven by diverse microbial mechanisms

Given the simulated dataset described above (for which uptake and secretion fluxes are known), we applied our contribution framework to calculate the contribution of each species to the variation observed in each of the 52 variable metabolites (Figure S5). The resulting contribution values can be used as ground-truth information about the link between microbial activity and environmental metabolites.

To highlight the nature and utility of such contribution values, and to demonstrate how metabolic fluxes translate into contribution profiles, we first describe our results for several example metabolites (Figure 2). Putrescine, an amino acid fermentation product, is an example of the simplest case, in which one microbial species – *E. coli* – synthesizes a metabolite that is not utilized or modified by other community members. Variation in the environmental concentration of putrescine was hence fully determined by the level of secretion from *E. coli*, which is therefore assigned a relative contribution of 1 (Figure 2B). Tetradecanoic acid, in contrast, was introduced (at a constant rate) via the nutrient inflow and utilized by the three *Bacteroides* species in the community to varying degree (primarily by *B. ovatus* and to a slightly lesser extent by *B. thetaiotaomicron*). The calculated contribution values successfully attributed variation in the environmental concentration of this metabolite to these three species, and correctly captured the difference in the magnitude between their effects (Figure 2C). Variation in uracil, another metabolite introduced via the nutrient inflow, was mainly driven by large shifts in its uptake by *B. ovatus*, but this effect is partially masked by *E. rectale*, which reduced its uptake when *B. ovatus’* flux was high and vice versa. Other species also utilized uracil, but at relatively similar levels across samples, and accordingly with relatively little impact on its variation. These complex patterns were all captured by the contribution profile obtained by our framework, with *B. ovatus* assigned a high positive contribution, *E. rectale* assigned an intermediate *negative* contribution, and other species assigned relatively negligible contribution values (Figure 2D). More complex species-metabolite relationships were also accurately and effectively summarized. Contribution values for acetate, for example, reflected the cross-feeding interactions that underlie variation in its concentration (Figure 2E). It was introduced to the shared environment by several species (primarily *C. symbiosum*), but most of its variation ultimately depended on the level of uptake by *E. rectale*. Finally, the contribution profile of succinate demonstrates how extremely strong interspecies interactions can produce contribution values much greater than the observed variance (Figure 2F). In the simulated data, this metabolite was synthesized by *B. hydrogenotrophica*, but was almost always fully utilized by other community members. The calculated contributions suggest that if the synthesis of succinate by *B. hydrogenotrophica* would not have been offset by uptake from other species, the variance in succinate concentration across samples would have been 71.7 times higher than is actually observed. (Note that the difference between positive and negative is always 1.)

Examining the complete set of variable metabolites and calculated contribution values revealed similar patterns of interactions (Figure S5). Specifically, as for the metabolites discussed above, negative contributions and/or contribution values greater than 1 were widespread. Nearly all metabolites (50 out of 52) had at least one species with a negative contribution value, and 36 had at least one species with a contribution value greater than 1. Of the 32 other metabolites with negative contributions, 29 were present in the nutrient inflow and their negative contributions result from competition between species for their uptake. This prevalence of negative and extreme values suggests that strong negative interspecies interactions have substantial impacts on metabolite concentrations, and that often, observed variation in a given metabolite’s concentration is the complex outcome of multiple species generating and offsetting much higher variation.

It is also important to note that while the average metabolic uptake/secretion flux of each species and the magnitude of its contribution to a given metabolite were generally significantly correlated (Spearman, *p* < 0.01 for 49 of the 52 metabolites), the species with the highest flux was often *not* the largest contributor to variation (26 of the 52 metabolites). Similarly, the variance in a species’ flux was significantly correlated with its contribution for 48 of the metabolites, but for 9 metabolites the species with the most variable flux was still not the largest contributor (due to differences in whether variable flux generated by one species is compensated by variation in the flux of another). These findings suggest that even if the magnitude and variation of species uptake and secretion fluxes across a set of microbiome samples are known (rather than just the abundances of species, which is the only measure usually assayed), metabolic interdependence between species would still make true contributor species challenging to identify.

Combined, the observations above highlight the complex relationship between species activity and measured metabolite concentrations, demonstrating the important role of both direct and indirect species interactions. This complex relationship, observed even in the idealized settings of our simulation model, is potentially markedly more complex than what is assumed by many microbiome-metabolite association-based analyses.

### Correlation analysis fails to detect true microbial contributors to metabolite variation

Given our observations above, we next set out to comprehensively assess how well pairwise correlation analysis (commonly used for analyzing microbiome-metabolome data) can detect true taxonomic contributors to metabolite variance. Put differently, we evaluated the extent to which a correlation between species abundance and metabolite concentration across samples captures the true causative contribution of a species’ metabolic activity to observed metabolite variation.

Following numerous microbiome-metabolome studies (14, 23, 28, 51), we considered identifying species-metabolite relationships as a classification task, aiming to identify for each metabolite the set of species that are primarily responsible for the variation observed in its concentration across samples. To this end, we defined *key contributor* species for each metabolite as those with a contribution value greater than 10% of the total positive contribution values. This resulted in a set of 83 species-metabolite key contributor pairs, representing true links between species activity and metabolite variation. On average, each metabolite had only 1.6 contributors (Figure S6), although 7.5 species on average had utilized or synthesized each metabolite at any point. 31.3% of these contributions occurred via synthesis reactions, 66.3% via utilization, and 2.4% (2 instances) via both processes. We then calculated the Spearman rank correlations between species abundances and metabolite concentrations across samples, and used a p-value threshold of 0.01 to define significant correlation between species and metabolites. This produced a set of 191 significant species-metabolite correlations, representing putative species-metabolite links. Scatter plots of these species-metabolite abundance relationships are shown for several example pairs in Figure S7.

Comparing this set of significant species-metabolite correlations to the set of species-metabolite key contributors clearly illustrated the difficulty of using univariate associations to infer mechanistic contributions (Figure 3). Indeed, of the 191 significant species-metabolite correlations, the vast majority (141) were false positives (corresponding to a positive predictive value of only 26.2%), and did not represent true contributor relationships (Figure 3A). Moreover, more than a third of these false positive species-metabolite pairs (51 out of 141) had *no* mechanistic connection; i.e., the species did not ever use or produce the metabolite in question. Furthermore, for 12 variable metabolites (out of 52), none of the key contributors were successfully detected by a correlation analysis. The overall accuracy was somewhat higher (66.5%), reflecting the high number of non-contributors that are also not correlated. Using a stricter cutoff (*p* < 0.0001, equivalent to a Bonferroni-corrected value of 0.05) only improved the positive predictive value to 33% and the accuracy to 77.1%. Indeed, a ROC curve analysis (Figure 3B) produced an area under the curve of 0.72, and overall correlations and scaled contribution values were only weakly associated (Figure 3C), suggesting that these findings can only be partially mitigated by changing classification thresholds. Metabolites of different classes had generally similar correspondence between correlations and contributions (Figure 3D).

**Figure 3.**
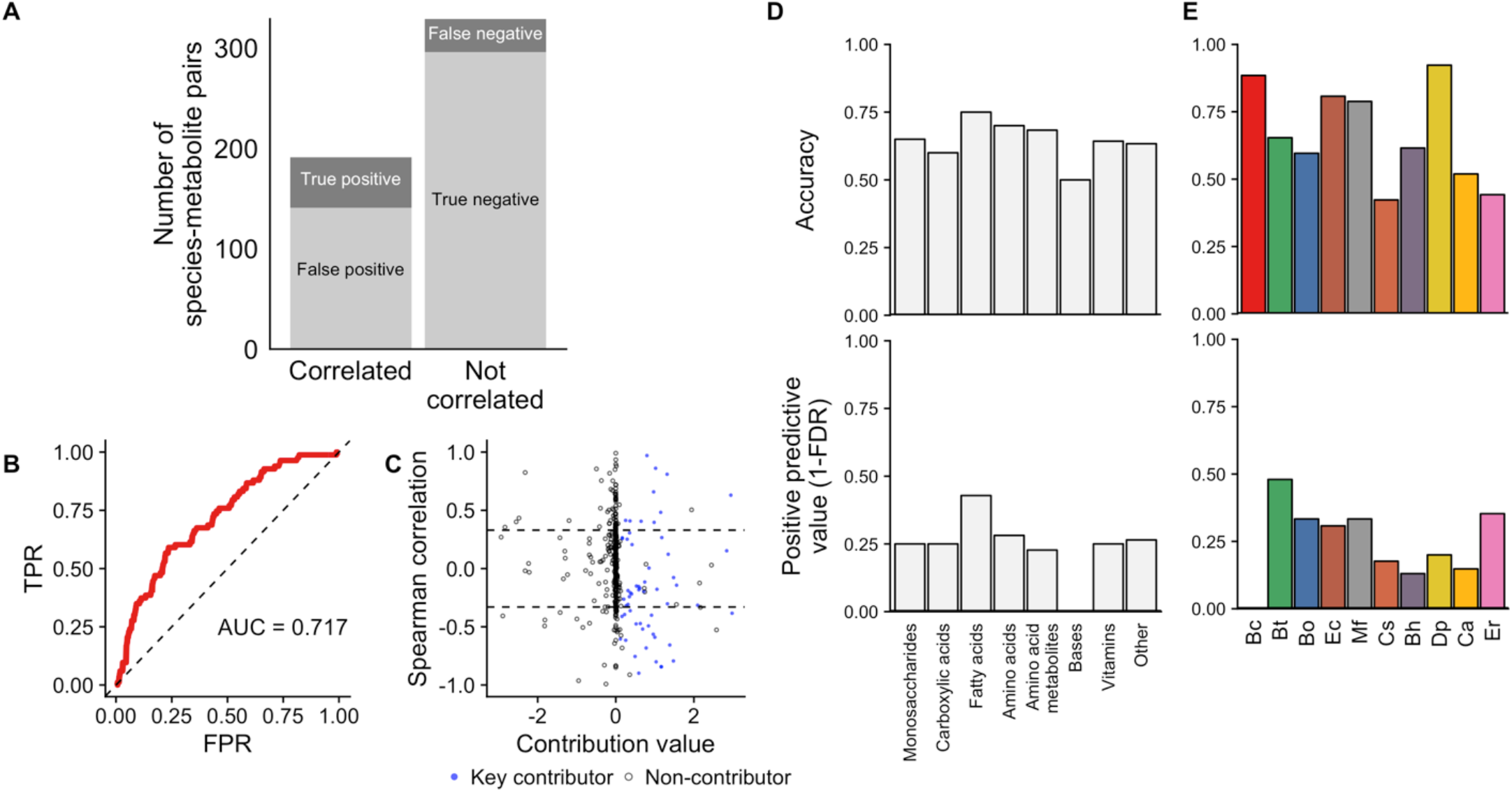
Species-metabolite correlations poorly predict species contributions to metabolite variation. **(A)** The number of species-metabolites pairs that were significantly correlated (left bar) or not-correlated (right bar) and its correspondence with true species-metabolite key contributors (indicated by shade of gray). **(B)** Receiver operating characteristic (ROC) plot, showing the ability of absolute Spearman correlation values to classify key contributors among all species-metabolite pairs. **(C)** Scatter plot of species-metabolite pairs, showing the poor correspondence between true contribution values (x-axis) and Spearman correlation (y-axis). Key contributors are plotted as blue points, others as hollow circles. Dashed lines show significant correlations (*p*<0.01). There are 65 species-metabolite pairs with a contribution value greater than 3 in magnitude whose values are not shown. **(D-E)** Accuracy and positive predictive value of Spearman correlation analysis for detecting true key contributors across metabolite classes (Panel D) and for each of the 10 species (Panel E).

Notably, key contributors for purely microbially-produced metabolites were not identified more accurately than those for metabolites in the nutrient inflow (66% versus 67%), which is perhaps not surprising since we used a constant inflow across samples (but see also our analysis below with variable inflow). Moreover, the total variance in a metabolite was not associated with the accuracy or predictive value with which key contributors for that metabolite were identified (Spearman rho, *p* > 0.1). Across species, contributions were identified most accurately for *D. piger*, which had a relatively low number of contributions (Figures 3E and S5C), but the positive predictive value was nonetheless <50% for all species.

We obtained similar results across several variants of this analysis (Supplementary Results, Figures S6, S8, and S9). To assess the impact of dynamic shifts over the duration of each simulation, we calculated an alternative set of contribution values based on the net steady-state metabolite flux rates at the final time point of each simulation, finding extremely similar results as for contributions to cumulative variation in concentration. We also evaluated the use of an alternative classification task, aiming to detect all microbes that affect variation in a given metabolite across samples regardless of whether their effects are ultimately reflected in the observed concentrations (i.e. those with large positive or negative contributions), again resulting in similar findings (Supplemental Results, Figure S6). Finally, we profiled the effects of model simulation parameters on correlation results, including the simulation length and the maximum enzymatic rate *V_max*, again finding minimal effects on contribution and correlation results (Supplementary Results, Figures S8-9).

### Species and metabolite properties explain discrepancies between correlations and contributions

Our analysis above demonstrated that correlations between species abundances and metabolite concentrations can often be only poorly associated with true contribution of species to metabolite variation. We therefore next investigated the origins of such discrepancies. We examined whether individual metabolites or species are predisposed to produce a significant species-metabolite correlation when the species in fact does *not* contribute to that metabolite variation (i.e., false positives), or to mask such correlation when the species *does* in fact contribute to this metabolite variation (i.e., false negatives), and if so, what species and metabolite properties are linked to those outcomes.

To determine whether the identity of the species or metabolite in question can explain inaccurate identifications of key contributors, we used a regression-based analysis. Specifically, we considered all species-metabolite non-contributor pairs, and fitted a logistic regression model to predict whether a species-metabolite pair exhibited significant correlation (false positive), based on either species identities, metabolite identities, or both (Methods). We then compared these three models using a likelihood ratio test to assess whether species and/or metabolite identities are informative. We similarly considered all species-metabolite key contributor pairs separately, again fitting a logistic regression model based on species identities, metabolite identities, or both to predict whether a pair failed to exhibit significant correlation (false negative).

For non-contributors, we found that false positives can be explained largely by species identity (likelihood ratio test (LRT) for inclusion of species terms *p* < 10^-13^). Incorporating both species and metabolite identities did not significantly improve the model (LRT for metabolite terms *p*=0.72). This finding suggests that false positives – correlations observed between species and metabolites to which they in fact did not contribute – are the outcome of interactions at the species level, regardless of the metabolite in question. This impact of strong interactions between dataset features on association test results has been described extensively in other data types (33, 34). Indeed, examining the 141 false positives identified above, we found that many can be explained by the relationships between the three dominant species in this community: *E. rectale, B. thetaiotaomicron*, and *B. ovatus*. These species competed strongly for carbon sources (and utilized their maximum allocation of sucrose, glucose, and fructose at nearly every step of the simulation), and their abundances were therefore negatively correlated. As a result, metabolites that varied due to the activity of one of these species were also frequently correlated with the other two. In total, 32 false positive correlations paired one of these species with a metabolite for which another species in this trio was a key contributor. More generally, we found that the probability of a false positive correlation for a particular species and metabolite depended on the species’ correlation with the true key contributors for that metabolite (*p*=0.006, Spearman rho between share of false positives and interspecies correlation; Figure 4A). Moreover, the maximum correlation each species had with any other species is a strong predictor of its overall specificity, which varies widely from 33.3% for *E. rectale* to 92% for *D. piger* (Spearman rho=-0.84, *p*=0.002). We also found that species identity was similarly predictive of whether a significantly correlated metabolite-species pair represented a true contributor versus a false positive (Supplementary Results).

**Figure 4.**
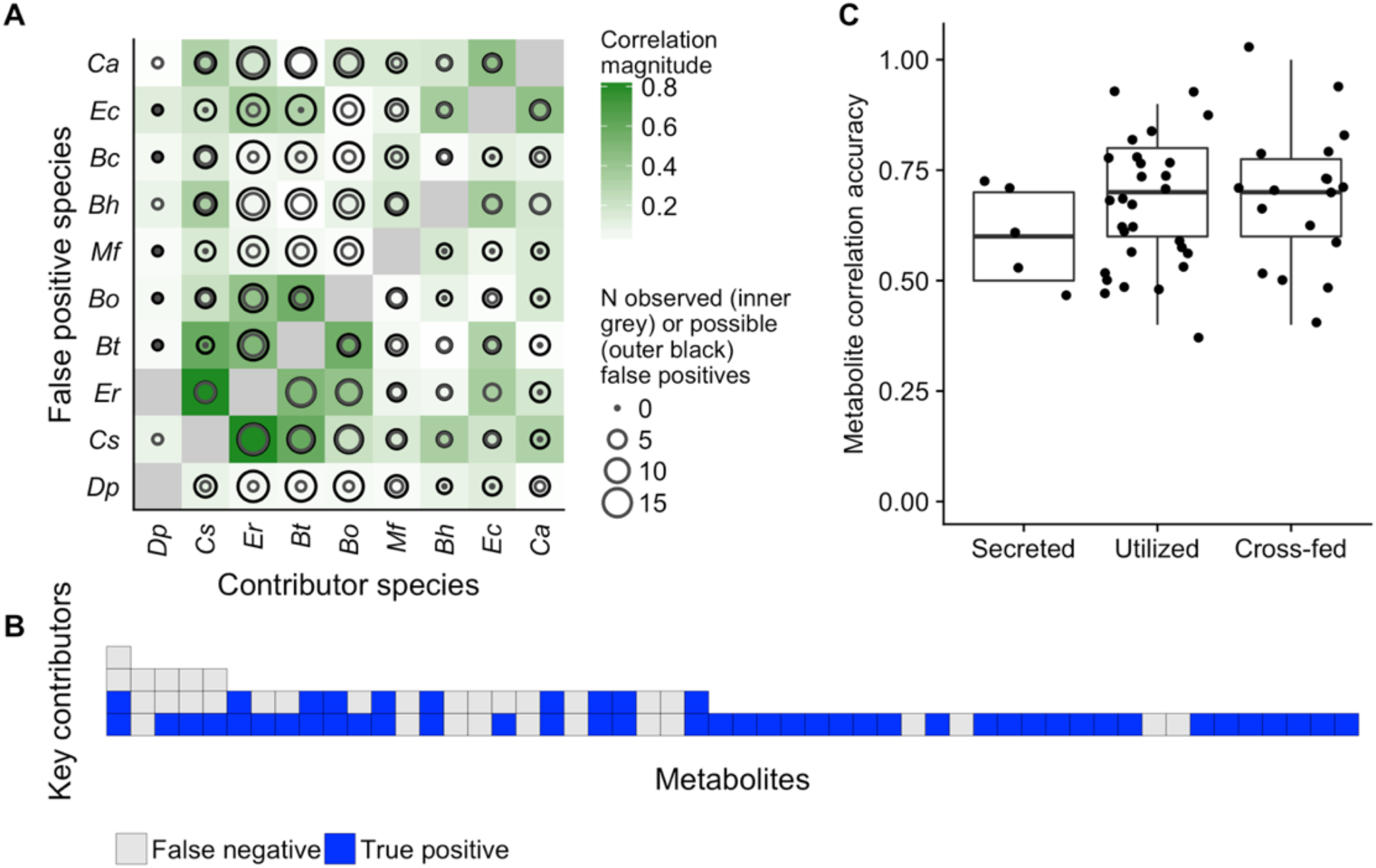
Metabolite and species properties explain correlation-contribution discrepancies. **(A)** Strongly correlated species pairs produced more false positive metabolite correlations. In this plot, the color of each tile indicates the strength of correlation in the abundances of each pair of species. The size of the outer black circle in each cell represents the number of metabolites for which the species on the x-axis is a key contributor and the species on the y-axis is not. The size of the inner circle represents the share of those metabolites for which a false positive is observed for the species on the y-axis. It can be seen that many false positive correlations involve the taxa with the strongest interspecies associations: *E. rectale, B. ovatus*, and *B. thetaiotaomicron*. **(B)** Metabolites with more microbial key contributors were more prone to false negative correlations. Each column represents an analyzed metabolite, ordered by its number of key microbial contributors, which are represented by each tile. The tiles are coded by the correlation outcome for each contributor. **(C)** Correlations detected key contributors equally accurately regardless of whether a metabolite is secreted, utilized, or cross-fed by the species. Each point represents the accuracy of correlations for a single metabolite across its comparisons with all 10 species.

In the case of key contributors, we found that false negative correlations can be explained largely by metabolite identity (LRT for metabolite terms *p*=0.002; although the species involved was also somewhat informative with LRT *p*=0.08). Put differently, a lack of correlation between the abundance of a key contributor species and the concentration of the metabolite to which it contributed was determined mainly by the nature of the metabolite in question. This lack of correlation between a given metabolite and its contributors could have resulted from competition or exchange of a metabolite between multiple species, such that none of the involved species end up strongly associated with the final outcome on their own. Indeed, across all metabolites, the average correlation between a metabolite and its key contributors is negatively associated with its number of key contributors (Spearman rho=-0.45, *p*=0.0008). The number of key contributors for any metabolite was also thus negatively associated with the sensitivity of contributor detection for that metabolite (Spearman rho=-0.48, *p*=0.0004; Figure 4B). We further hypothesized that false negative outcomes might be more common for metabolites with more or larger negative species contributions, since these, by definition, mask or compensate for the activity of key contributor species. While all metabolites with a false negative outcome did have at least one species with a negative contribution value, as mentioned above, this was true for nearly all analyzed metabolites (50/52), and the number of negative contributing species was not associated with the occurrence of a false negative correlation (*p*=0.86, Wilcoxon rank sum test). Moreover, we also did not observe any effect of the average concentration of a metabolite on the sensitivity and accuracy of its detection via correlation analysis, nor of whether it is secreted, utilized, or cross-fed (Figure 4C). In summary, our analysis suggests that the largest factor explaining whether a metabolite’s key contributor can be detected by a correlation analysis is simply whether there are other community members (key contributors) that also impact the observed concentration of that metabolite.

### Environmental fluctuations in metabolite concentrations impact detection of key contributors

Our analyses above all focused on a single simulated dataset in which the nutrient inflow was constant across all samples, meaning that metabolite variation was fully governed by microbial activity. However, in reality, metabolite variation can and does arise also from non-microbial sources, potentially affecting both the landscape of key microbial contributors and our ability to detect them via correlation-based analyses. To explore the impact of environmental fluctuations, we therefore ran several sets of additional simulations with varying degrees of nutrient fluctuation, designed to emulate a range of levels of experimental diet control and variation in host absorption across the simulated mouse gut communities. In these simulations, we maintained the same set of 61 initial species compositions but added small amounts of stochastic noise to the nutrient inflow, sampling inflow concentrations for each compound in each simulation from a normal distribution with a mean equal to the compound’s original inflow rate and a standard deviation ranging from 0.5% to 10% of the mean in 8 increments (Methods). For each of the resulting 8 datasets, we again calculated contribution values (with the added element of the nutrient inflow as a potential contributor to variance), identified key contributors, and compared them with the results of a correlation analysis.

Examining the obtained contribution values, we found, as expected, that variation in inflow quantities can outweigh the variation in microbial fluxes, and that as the variation in inflow increases, its contribution to metabolite variation increased at the expense of the contributions of community members (Figure 5A). As a result, the number of key contributions attributed to each species decreased for metabolites in the nutrient inflow (Figure 5B). Interestingly, however, some species lost their contributions more gradually than others, and in some cases even became key contributors for additional metabolites (Figure 5B). For most metabolites, the relative ranking of species with the highest contribution values was unchanged with increasing fluctuations (Supplementary Results).

**Figure 5.**
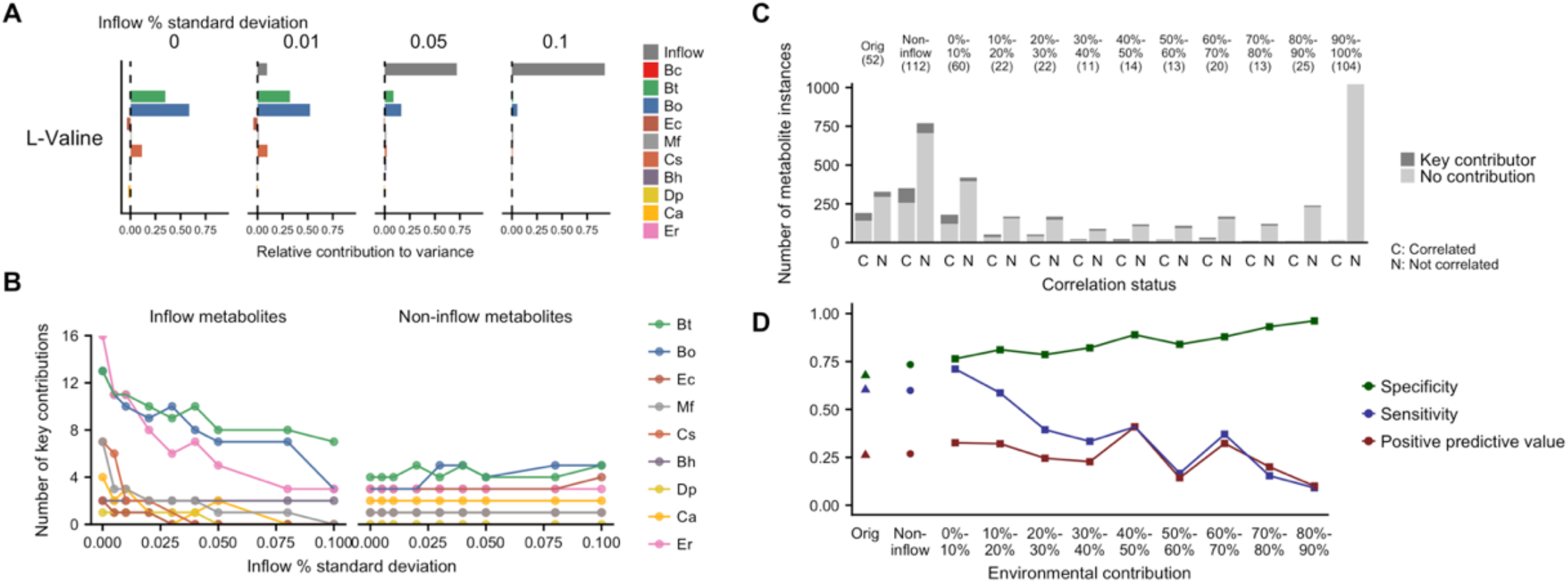
Environmental fluctuations impact correlation-contributor sensitivity and specificity. **(A)** Example set of contribution profiles for a single inflow metabolite, L-valine, with increasing fluctuations in its inflow. The relative contribution values for each species and for the inflow are shown for 4 sets simulation runs, each with a different degree of fluctuation. The label on each plot describes the relative standard deviation (coefficient of variation) of inflow metabolite concentrations for that set of simulations. The microbial contributions to variance in L-valine concentrations became relatively smaller with increasing variation from the external environment. **(B)** Shifts in key microbial contributors with increasing environmental inflow fluctuations. The number of key contributions of each species to the 52 analyzed metabolites is shown, separately for metabolites present in and absent from the nutrient inflow. Microbial contributors to inflow metabolites decreased as environmental contributions increased, but this effect varied between taxa. **(C)** Correlation analysis failed to detect key microbial contributors regardless of the size of contribution from external inflow variation. Across all sets of simulations, metabolites were binned based on the percent of total positive contribution from the external inflow. The bar plots shown have the same format as Figure 3A, showing the number of species-metabolites pairs that were significantly correlated (left bar) or not-correlated (right bar) and its correspondence with true species-metabolite key contributors (indicated by shade of gray). The first two bars, labeled “Orig” describe the original set of simulations (replicating Figure 3A). The next two show the results for non-inflow metabolites across all levels of inflow fluctuations. The remaining bars show the results for metabolites with increasing levels of environmental contribution. **(D)** Correlation analysis detected key microbial contributors with increased specificity, decreased sensitivity, and generally consistent positive predictive value with increasing contribution from the external inflow. Sensitivity, specificity, and positive predictive value are shown for same environmental contribution bins as in Panel C.

We next examined how correlation-based detection of key microbial contributors was affected by these inflow fluctuations. We assigned each of the 52 metabolites in each of the 9 datasets (the original dataset with no inflow fluctuations and the 8 datasets with varying degree of fluctuations) to bins according to the level of contribution attributed to the inflow for this metabolite at that degree of fluctuation (see Methods). We then evaluated the performance of correlation analysis for each bin separately. The share of true key contributors naturally decreased rapidly with increasing environmental contribution, as did the number of significantly correlated species-metabolite pairs (Figure 5C). Importantly, however, the sensitivity of correlations decreased substantially with the level of contribution attributed to the inflow, but the specificity in fact increased from 67.7% to 92.3% (Figure 5D). This suggests that while environmental fluctuations disrupted the signal linking microbial species with the metabolites they impact, they also disrupted indirect associations between species and metabolites (false positives). Overall, however, the AUC did not change significantly with increasing environmental contribution (Figure S10A), and the positive predictive value is similarly relatively stable (and never rose higher than 37%). Interestingly, the detection of some metabolites not present in the inflow was also affected by inflow fluctuations in a similar manner (Supplementary Text, Figure S10B).

### Correlation analysis is similarly limited in simulations of more complex and diverse human gut microbiota

Our results have illustrated consistent discrepancies between microbe-metabolite correlations and microbial contributions to metabolite variation in a model ten-species community. We lastly addressed the question of whether these findings generalize to more complex mammalian gut microbiota, communities with many times more taxa and a more uneven distribution across individuals. To do so, we ran an additional set of simulations emulating human gut microbiota transplanted into gnotobiotic mice. We first mapped 16S rRNA sequence variants from the Human Microbiome Project (52) to the genomes of the AGORA model collection at 97% sequence identity (40), and selected 57 samples with a successful mapping rate greater than 25% relative abundance. The total share of mapped reads averaged 36.7% across these samples, with a maximum of 73.5%. Despite this variation, mapped reads displayed features typical of Western gut microbiomes, including a predominance of Bacteroidetes and Firmicutes phyla along with varying lower abundances of Actinobacteria and Proteobacteria (Figure 6A). The number of species identified in each sample ranged from 23 to 62, with a median of 42. We ran a simulation based on each sample by setting the initial species relative abundances according to the relative abundances of mapped reads, while maintaining the same physical parameters as previous simulations (see Methods for additional details). We used nutrient inflow quantities with 1% standard deviation between samples. Initial species compositions displayed characteristic shifts in abundance over the simulation time course (Figure S11A). Metabolites were also highly variable, with a median coefficient of variation of 71% across 222 metabolites (Figure S11B). We calculated contribution values for this dataset, finding a smaller share of key contributions (only 392 out of 29,082 possible species-metabolite pairs). Only 35.1% of species (46 out of 131) were identified as key contributors to any metabolite. The genera with the most contributions were *Bacteroides, Ruminococcus*, and *Enterobacter*, which were also three of the four most abundant genera in the final dataset (Figure 6B).

**Figure 6.**
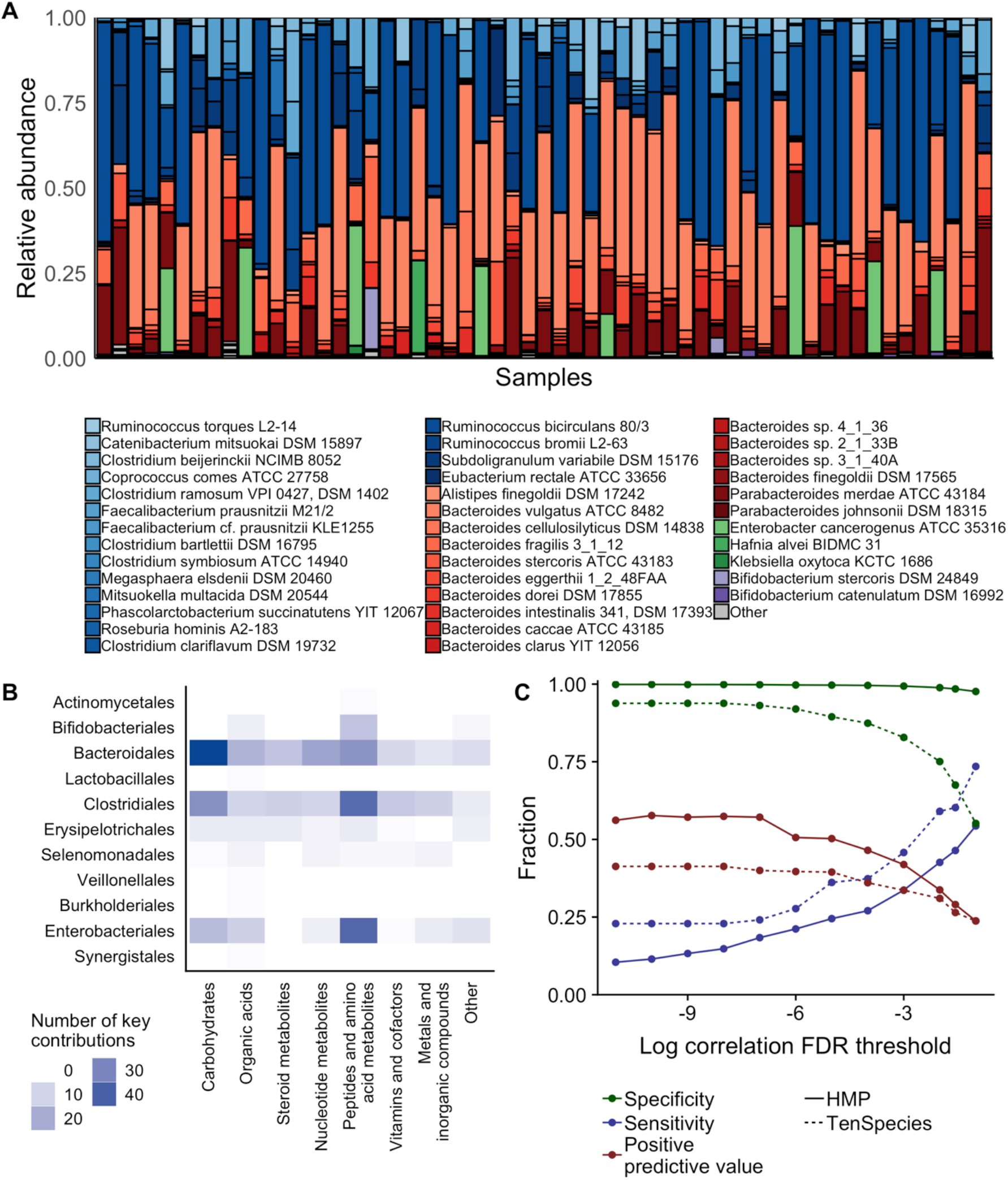
Correlation-contribution discrepancies persist in simulations of complex human gut-based microbiota. **(A)** Species abundances of the 57 Human Microbiome Project (HMP) based-simulations at the 144 hour time point. Shades of blue indicate species in the phylum Firmicutes; red, Bacteroidetes; green, Proteobacteria; and purple, Actinobacteria. **(B)** Key contributions to metabolite variation across the HMP-based dataset, summarized at the level of taxonomic orders and metabolite categories. **(C)** Performance of correlation analysis for identifying key species-metabolite contributors in the HMP-based dataset (solid lines) compared with the original 10-species dataset (dashed lines) across varying significance levels, using Benjamini-Hochberg false discovery rate (FDR) corrected *p*-values.

**Figure 7.**
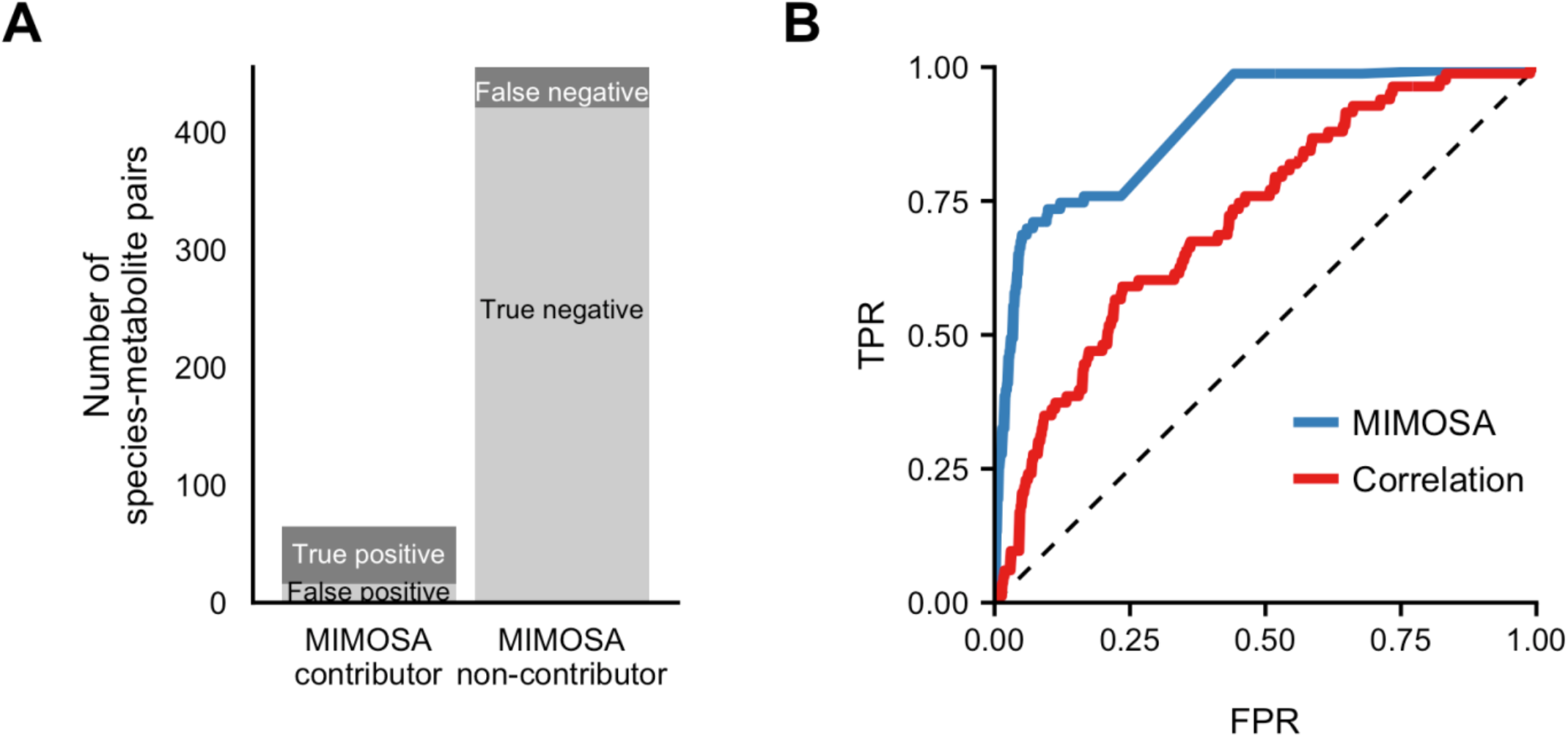
MIMOSA identified key microbial contributors more accurately than correlation analysis. **(A)** The number of species-metabolite pairs that were identified as potential contributors (left bar) or not (right bar) by MIMOSA, and its correspondence with true key contributors. **(B)** Receiver operating characteristic (ROC) plot, showing the ability of both MIMOSA and absolute Spearman correlation values to classify key contributors among all species-metabolite pairs.

In this noisier and more layered dataset, only a small share of species-metabolite pairs was significantly correlated. In order to fairly compare with the previous dataset while accounting for the larger number of hypothesis tests, we defined significance based on an equivalent Benjamini-Hochberg estimated false discovery rate (0.027) as the *p* < 0.01 cutoff used for the previous dataset. 2.2% of species-metabolite pairs displayed significant correlations at this cutoff (*p* < 0.00058). This level of correlation is comparable to a recent microbiome-metabolome study of the colon of healthy humans (51), in which 1.4% of OTU-metabolite pairs displayed Spearman correlation coefficients of the same effect size. In our dataset, correlation analysis detected contributors with high specificity (98.4%), and an area under the ROC curve of 0.89. However, the positive predictive value was still only 29.0%, rising as high as 57% with a significance cutoff of *p* < 10^-10^. We compared these classification results with the original dataset, finding that despite the difference in overall AUC, sensitivity, sensitivity, and predictive value are similar or worse for the two datasets at commonly used FDR thresholds between 0.1 and 0.01 (Figure 6C), and sensitivity and predictive value are both highly dependent on the choice of significance threshold. As in the ten-species dataset, a large share of false positive species-metabolite pairs (65.4%, 291 out of 445) also involved species with no capacity to impact the metabolite in question.

The outcomes of correlation analysis were influenced by the same factors as observed in the model community dataset, but also by several additional characteristics. False positive classifications were, again, driven by interspecies covariance: Species significantly correlated (at 10% FDR) with a true key contributor for a metabolite were 13.6 times more likely to have a false positive correlation with that metabolite than species with no such link (*p* < 10^-16^). Notably, the false positive rate of a given species was also substantially affected by its prevalence: the number of samples in which a species was present was negatively associated with its specificity (Spearman rho = −0.57, *p*=0.002, Figure S11C), among species with at least 3 key contributions. In other words, widely prevalent species were more prone to false positive correlations than rarer species.

False negative contributions were again influenced by properties of both metabolites and species. As in the ten-species dataset, species contributions to metabolites with more than one key contributor were 5.2 times more likely to not be correlated than those that were the sole key contribution for a metabolite (*p* < 10^-10^, Fisher exact test). In this dataset, an elevated share of these metabolites with multiple key contributors were cross-fed between different species (*p*=0.00007, Fisher exact test), and correspondingly, key contributors for cross-fed metabolites were also 1.6 times less likely to be significantly correlated (*p*=0.02). Both cross-feeding and false negative outcomes occur variably across metabolite classes, with nucleotide metabolites having the highest rates of both phenomena (Figure S11D). Taken overall, our simulations and analysis of this realistic microbiota simulation demonstrates that correlation analysis can have greater utility in a microbial community dataset with greater complexity and variability, but the results are again strongly influenced by properties of individual metabolites and species.

## Discussion: Insights and implications for microbiome-metabolome analyses

Above, we have investigated the ability of correlation-based analyses to detect key microbial contributors responsible for variation in metabolite concentrations across samples. Our findings suggest that microbe-metabolite correlation analysis may be a useful approach for exploratory analyses, but they highlight some of the limitations and caveats of such microbiome-metabolome studies and identify several factors that impact the relationship between community composition and metabolite concentrations. Below, we elaborate on a set of practical conclusions and their implications for the analysis and interpretation of microbiome-metabolome studies.

### Association-based analyses of microbiome-metabolome assays have low predictive value for detecting direct species-metabolite relationships and require conservative interpretation

Microbiome-metabolome association studies have been previously proposed as a powerful tool for the identification of causal mechanisms of microbiome metabolism (53), and indeed, such studies often present detected associations as evidence for mechanistic relationships (11, 27, 29). However, our analysis suggested that the positive predictive value of significant species-metabolite correlations for identifying true microbial contributors can be extremely low: less than 50% across all settings, as low as 10% in the context of large environmental fluctuations, and 29% in simulations based directly on human gut composition. Recent experimental studies pairing microbiome-metabolite correlation analysis with *in vitro* monoculture validations have similarly anecdotally observed many false positive correlations (32). Additionally, given the somewhat low sensitivity observed in our analysis, a lack of association is not necessarily sufficient to reject a hypothesis that a particular microbial taxon impacts a particular metabolite. The choice of correlation threshold should therefore be chosen carefully, taking into account the complexity of the community and the environmental context. In general, identified correlations between microbial taxa and metabolites should be interpreted very conservatively and used mostly to prioritize microbe-metabolite relationships for follow-up validation studies (e.g., via culture-based studies or germ-free model organism colonization). One potential approach for improving the predictive value of such correlation-based analyses is to examine whether they replicate across multiple conditions. Indeed, we found that a correlation does provide stronger evidence for a contributor relationship if it persists across different contexts. Across our 9 simulated datasets with varied environmental fluctuations, the 43 species-metabolite pairs that were significantly correlated in every dataset were 2.1 times more likely to denote true key contributor relationships than other significant correlations (Fisher exact test, *p*=0.05), although their positive predictive value was still relatively low (39.5%). Of the limited number of significant correlations shared between our original and HMP-based datasets (*n*=5), all were false positives in both datasets, reiterating the need for caution.

### The predictive power of correlation-based analysis is species-, metabolite-, and context-dependent

In our datasets, metabolites varied widely in both contribution profiles and in their detectability via correlation analysis. In particular, the key contributors for metabolites acted upon by fewer species, and potentially those that are not exchanged between different species, were identified more readily. Moreover, in our simulations of human gut communities, contributions by less prevalent species were identified much more accurately than those by widely-found species, indicating that hypotheses based on associations of rarer species should potentially be prioritized. Correlation analysis may thus identify microbes involved in specialized secondary metabolic processes (e.g. products of complex biosynthetic pathways) more readily than those involved in more widespread processes.. Therefore, correlation-based approaches may be more informative for analyzing compounds that are specific to a small number of rare taxa, but accurate dissection of the taxa controlling variation in widely-trafficked metabolites may require more detailed analysis and experimentation. Similarly, we found that species-metabolite correlations for species that are strongly associated with other taxa (e.g., those with tight interactions with other community members) are often spurious, suggesting that such correlations should be regarded less confidently.

### External metabolic fluctuations can strongly impact the detection of microbial contributions

Our analysis of the impact of environmental fluctuations suggested that the presence of environmental variability from a diverse set of samples could in fact increase correlation specificity. We also found that the sensitivity of correlation analysis rapidly decreased with increasing environmental fluctuations (from 60% to 9%). These observations suggest that while a tightly controlled environment (e.g., identical diets) is intuitively expected to increase the strength of microbiome-metabolome studies, its value depends on the study priorities. Specifically, if the goal is to identify clear-cut microbial drivers of healthy- and disease-associated metabolite shifts, stochastic variation in nutrient availability could be beneficial as it may reduce the rate of false positive associations. In contrast, for studies searching for a particular microbial taxon’s involvement in a particular process (e.g. aiming to determine whether an ingested probiotic impacts aspects of gut metabolism), a more controlled environment may be favorable. It should, however, be noted that our findings were based on environmental fluctuations that were uniform and independent, which may not hold for real-life environmental fluctuations such as diet variation. It is also worth noting that in our simulations, microbial fluxes for some environmental metabolites could be drowned out by as little as 0.5% variation in nutrient inflow quantities, while others still had substantial microbial contributions even with 10% variation in inflow. When interpreting an observed association, the scale of possible microbial variation relative to external variation should therefore be taken into account.

### Mechanistic reference information can improve the predictive power of microbiome-metabolome studies

In our simulated dataset, 36% of the false positive correlations occurred between a metabolite and a species that was in fact not capable of uptaking or secreting that metabolite. Ruling out such falsely detected links would substantially improve the positive predictive value of a correlation-based analysis. One approach for doing so is by utilizing genomic information, which can be obtained or predicted for many microbial taxa (54). By coupling such genomic information with metabolic databases such as KEGG or MetaCyc (55, 56), researchers can filter out correlation-based links that are likely not feasible causative relationships. Further improvement can be obtained by integrating such reference information directly into the analysis. Indeed, we previously introduced a computational framework, termed MIMOSA (57), that utilizes a simple community-wide metabolic model to assess whether measured metabolite variation is consistent with shifts in community metabolic potential, and to identify potential contributing taxa. MIMOSA has been applied to varied host-associated microbiomes from varied body sites and from human and mouse hosts (12, 58, 59). Applying MIMOSA to the simulated ten-species dataset analyzed above (Methods), we found that it indeed identified key contributors significantly more accurately than a correlation-based analysis, with an AUC of 0.89 (Figure 6). Notably, in this analysis, we assumed MIMOSA has access to the correct set of metabolic reactions possessed by each species. Using standard less-complete information obtained directly from the KEGG database (as done regularly when using this tool) reduced the number of metabolites that could be analyzed from 52 to 39, with improved specificity (96%) and positive predictive value (61%) and an ultimately comparable AUC (0.74). Combined, these findings suggest that reference model-based approaches can provide stronger evidence for mechanistic relationships than strictly correlation-based methods, but their use depends on complete and high-quality metabolic reference databases.

## Future opportunities and challenges

Microbiome-metabolome studies have an important role in microbial ecology research. They specifically have great potential to dissect the metabolic interactions of complex microbial communities, and to unify “top down” and “bottom up” microbiome research approaches by providing mechanistic information at a systems level. Moreover, from a translational perspective, microbiome-metabolome studies can inform efforts to design targeted therapies to alter specific microbial or metabolic features of a community (13). Such interventions require first identifying putative targets, which in many cases may entail identifying the key contributor species that drive observed shifts in a particular beneficial or detrimental metabolic phenotype.

Importantly, while we show here that a correlation-based analysis may be limited in its ability to identify these key microbe-metabolite links, this does not necessarily imply an inherent limitation of microbiome-metabolome data. For example, analyzing our data, we found that species abundance is in fact a very good proxy for metabolic activity (median correlation of 0.996 between abundance and flux for all species-metabolite pairs), meaning that the variance in total species abundance drastically outweighs the individual-level variance in flux rates. When we further examined whether false negative associations in our original dataset stem from a disconnect between the abundance of a species and its metabolite uptake or secretion rates, we identified only 2 undetected key contributor pairs that could be explained by such a discrepancy. This analysis suggests that taxonomic abundance data is sufficient to explain and model community metabolic variation to great extent, despite common concerns about potential discrepancies between community composition and function. It also suggests that metatranscriptomic expression data may not provide much additional value for this purpose, as other studies have indicated (54, 60, 61).

Given the increasing prevalence of microbiome-metabolome studies, their promise, and the caveats of association-based research discussed above, further development of computational and statistical methods for analyzing such datasets is clearly needed. Possible directions include the use of multi-species dynamic metabolic models that can replicate experimental observations (62), multivariate approaches for deconvolving interactions between species and the environment (63, 64), and probabilistic methods that can integrate prior information while allowing for other unknown mechanisms (31, 65). The conceptual framework of taxon-metabolite contributions, and the use of dynamic simulations demonstrated here, can both inform the future development and evaluation of such methods.

There is also a continued need for gold standards to evaluate new methods. This study is only a first step in that direction and has analyzed one specific type of research question: identifying microbial taxa directly responsible for variation in metabolite concentrations between samples in a cross-sectional study design. Although this focus describes many recent microbiome-metabolome studies, other studies may address a wide range of complementary research questions, and correspondingly, the desired “ground truth” can take different forms. Moreover, depending on the objective, an alternative definition of a taxon-metabolite relationship may be required. For example, it may be valuable to identify key contributors that act via alternative mechanisms, such as by modifying substrate availability or environmental conditions (for example (66)), or to distinguish metabolite variation arising in response to a perturbation from variation due to differences in steady-state metabolism between communities. Additionally, our findings rely on an *in silico* system that may not capture many aspects of community ecology and metabolism, and it is possible that the predictive value of correlation analysis, as well as of other analytical methods, differs fundamentally in this system as compared to true biological systems. Further studies should also consider additional variables such as community diversity, sample size, measurement error, and other types of environmental variation. Ongoing technology developments in mass spectrometry and stable isotope probing will ideally enable future evaluation analyses using experimental, quantitative, species-specific community flux data to define key microbial contributors (67, 68). Such evaluations can also take advantage of datasets comparing community microbiome-metabolome data with *in vitro* monoculture or mono-colonization data (32, 35, 36).

Ultimately, much remains to be learned about the many processes through which complex microbial communities shape their environment. The first major call for the application of metabolomics to microbiome research, published 10 years ago (69), noted that new methods will be necessary to integrate genomic and metabolic data and inform the prediction of community metabolic properties from metagenomes. Now that microbiome-metabolome datasets are widely available, ongoing development of analysis methods for these studies has great potential to generate new knowledge. Moreover, future work in this area stands to benefit from the utility of dynamic, multiscale metabolic modeling. Detailed mechanistic simulations are used widely in astronomy, climate science, and other fields to make methodological choices and assess possible experimental outcomes when ground truth measurements are unavailable or difficult to obtain (70, 71). An analogous strategy in microbiome research may be similarly fruitful.

## Methods

### Derivation of species contributors to variation

We derived an expression representing the contribution of each species to the variance in the concentration of each metabolite. While we describe this calculation in terms of species, a similar calculation could be done at the level of phyla, strains, or any grouping of the community for which metabolite secretion and uptake fluxes are available.

The concentration of a given metabolite *M* at the end of a single simulation run is a function of the uptake and secretion fluxes (responding to the species’ degradation and synthesis activities) of the *n* species, the environmental inflow over all time steps *m_in_*, and the dilution *m_out_* out of the chemostat over all time steps:

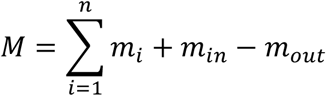

The value of *m_out_* at a given time step *t* is the product of the dilution rate *D* and the metabolite concentration at the previous time point (see above). This fact can be used to express *m_out_* in terms of all the previously recorded environmental inflow and microbial activities. The metabolite concentration at any time point *t*, *M(t)*, is then equal to:

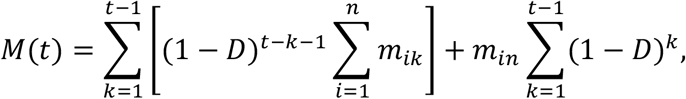

where *m_ik_* represents the activity of species *i* at a single time point *k*. We can then ignore dilution outflow by replacing each activity value *m_i_* in the final concentration calculation above with a value corrected for the mitigating effect of chemostat dilution over the course of the simulation up to time *t*, defined here as *m_i_^*^*. *m_i_^*^* represents the total amount of a compound secreted or uptaken by species *i*, minus the share of that quantity that is eventually diluted out over the course of the simulation.

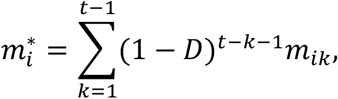

and thus,

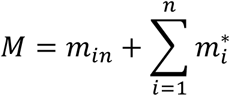

In this work, we refer to “environmental fluctuations” as the effect of the independently parameterized nutrient inflow, *m_in_*, and where not otherwise specified we use *m_i_* to imply *m_i_^*^*, a species activity quantity that accounts for the corresponding subsequent dilution out of the system.

Using the expression above, *var*(*M*) can then be clearly expressed as a sum of correlated environmental and microbial random variables:

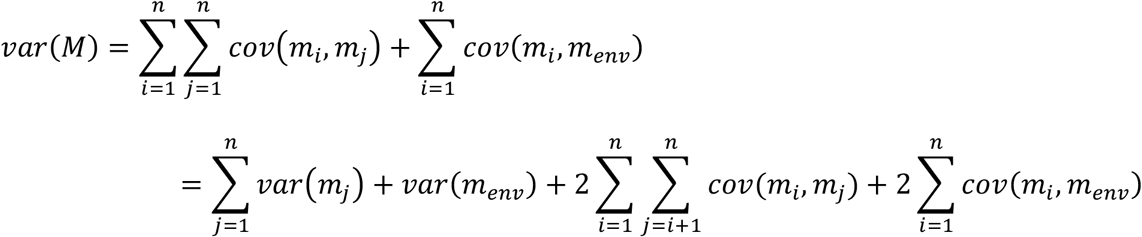

This expression can then be partitioned additively into *n*+*1* terms representing the contribution of each microbial species and of fluctuations in the environmental nutrient inflow.

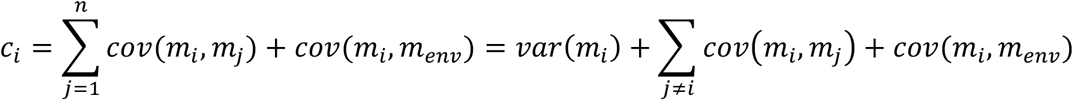

### Multi-species Dynamic Flux Balance Analysis modeling

In this study, we simulated the growth and metabolism of a community of 10 representative gut species that was previously explored experimentally (47). We specifically utilized a previously introduced multi-scale framework for modeling the dynamics and metabolism of multiple microbial species in a well-mixed shared nutrient environment (44, 72). This framework assumes that each species in the community aims to maximize its own growth on a short time scale given available nutrients, and uses Flux Balance Analysis to predict the growth and metabolic activity of each species at this short time scale (46). The shared environment is then iteratively updated based on the species’ predicted growth, uptake, and secretion rates, such that metabolic interactions are mediated via the environment as a natural byproduct of species activities, rather than being explicitly modeled (45).

We used genome-scale metabolic model reconstructions of the 10 community members from the AGORA collection version 1.01 (40), which have been consistently curated to remove or modify thermodynamically unfavorable reactions, remove futile cycles, and confirm growth in anaerobic environments on expected carbon sources, with additional curation for several biosynthesis pathways. The COBRA toolbox was used to convert each AGORA model to MATLAB format (73). The growth and metabolism of the 10-species community were simulated in a chemostat setting in 15-minute time intervals. We set the chemostat volume to be approximately equal to a mouse gut (0.00134 liter (74)). We similarly set metabolite inflows to emulate the macronutrient and micronutrient quantities in a corn-based mouse chow (47) (provided in Supplementary Data 1).

The simulations were performed following a previously introduced procedure (44), repeated for each time step *t_n_*: First, the maximum uptake rate for all metabolites by all species, denoted as *ν_jk_* for metabolite *j* and species *k*, were calculated based on Michaelis-Menten single-substrate kinetics, with assumed universal values for maximum rate *V_max_* and transporter affinity *K_m_* for all metabolites (provided in Supplementary Data 1). *ν_jk_* was further constrained based on an allocation of the metabolite’s environmental concentration to each species in proportion with its biomass. Then, the steady state reaction fluxes for each species *k* at time point *t_n_* were determined by maximizing the growth rate *μ_k_*, within the obtained constraints on environmental metabolite uptake. To obtain a single and consistent flux solution for each species, the total flux activity for each species (i.e., the sum of absolute fluxes given the predicted optimal growth rate) was minimized, under the assumption that organisms prefer to operate their metabolism with minimal enzymatic cost (75). The optimal flux solutions were solved using linear programming with GLPK (www.gnu.org/software/glpk). With the resulting flux and growth rate information, the total biomass of each species *k*, *bio_k_*(*t_n_*), was updated for the next time point *t_n+1_*, using a standard exponential growth function incorporating dilution:

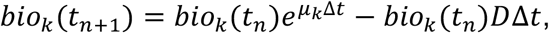

where *D* is the dilution rate. We set *D* to 0.0472 per hour, in order to obtain community growth rates consistent with the observed average growth rate of the three most abundant species growing under 47 different carbon conditions (76). The total amount of uptake or secretion for each species *k* and metabolite *j* over a single time step was then calculated as previously derived (44):

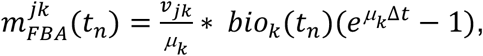

where *ν_jk_* is the rate of uptake or secretion specified by the FBA solution for that species and metabolite at that time point, *μ_k_* is the species growth rate, *bio_k_*(*t_n_*) is the species abundance, and Δ*t* is the size of the time step. Finally, combining the flux solutions of all species, nutrient inflow, and dilution, along with the steady state assumption of no intracellular metabolite accumulation, the concentration of a given metabolite in the shared nutrient environment at the next time point, *M_j_*(*t_n+1_*) can be updated as:

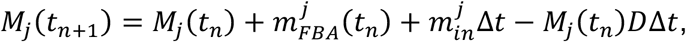

where 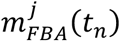 is the metabolic impact from all species considering their abundance and their uptake and secretion rates of metabolite *j*, and 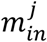 is the inflow rate of metabolite *j*. This process of calculating uptake rates, Flux Balance Analysis solutions, and updated metabolite concentrations was then repeated iteratively for the duration of the simulation.

Each simulation was run for a period of 144 hours or 576 time steps. This time period was long enough for most simulation runs to approach a steady state composition: specifically, in >65% of the simulations analyzed in our study, the change in abundance in any species over the final 3 hours was less than 0.01% of the carrying capacity (see below), and all had no changes greater than 0.3% of the capacity over that period. The concentrations of species and metabolites, the species growth rates, and the solved rates of all reactions for each species (including uptake and secretion) were recorded in each step of each simulation and used for subsequent analyses (Supplementary Data 1 and 2).

### Simulation initialization parameters

We fixed the initial total abundances of microbes to the carrying capacity for this system and media, which was estimated to be 0.433 units of biomass. This capacity was calculated as the average final total abundance from a set of simulations with varying compositions and low initial abundances. We then varied the relative abundances, increasing the abundance of one species at a time at the expense of all other species equally. Specifically, for each species, we ran simulations in which the ratio of that species’ initial abundance relative to all other species was 2, 3, 4.5, 6, 9, and 13 times (equating to a range in relative abundance of 10% to 60% for each species). This resulted in a total of 61 simulation runs (one with all species starting at equal abundance and 6 with increased abundance of each species). We chose this sample size to approximately represent the sample sizes of published cross-sectional microbiome-metabolome association studies (14, 16). We set the initial inflow concentrations to the amount that would dilute in over one hour under the calculated inflow rates.

### Calculation of contribution values for variable metabolites

We calculated contribution values for all metabolites with variance in concentration above the 25^th^ percentile. We chose this threshold in order to include as many metabolites as possible while excluding those that only varied at all in fewer than half of the simulation runs, or whose variation would be subject to potential numerical errors.

### Comparison with Shapley values

We implemented an approximate Shapley value algorithm (43) as an alternative strategy to calculate contributions for the simulated dataset. Briefly, 15,000 random orderings of the 10 species were randomly generated. For each ordering, the variance in metabolite activity is calculated for subsets of size 1 to 10, adding in species according to the specified ordering. The difference in variance as a given species is added to the subset, denoting the *marginal* contribution of that species to variation, is recorded. The average marginal contribution across all orderings for each species is then defined as its contribution to variance.

### Species-metabolite correlation analysis

We calculated Spearman correlations between absolute species abundances (quantified as total biomass) and concentrations of variable metabolites. We used absolute abundances in order to evaluate the relationships between species and metabolites under the hypothetically best possible measurements of both data types. We also compared correlation results using relative abundances and found very minimal differences in the main simulation dataset: only 7 species-metabolite pairs (1.3%) are significantly correlated using absolute abundances but not relative, and only 4 pairs (0.8%) are correlated using relative abundances but not absolute.

We used a *p*-value threshold of 0.01 to classify “significant” associations for binary comparisons. For interpretability, we refer to p-values not corrected for multiple hypothesis testing, since the number of tests remained constant across nearly all of our analyses (520 possible species-metabolite pairs). The 0.01 threshold we use to define significantly correlated pairs is equivalent to a Benjamini-Hochberg corrected false discovery threshold of 0.027, calculated using the R function *p.adjust* (77).

### Logistic regression modeling of correlation outcomes

We used logistic regression models to identify factors that can be used to predict whether a non-contributing species-metabolite pair displays a significant correlation (false positive), and whether a key contributor species-metabolite pair fails to be correlated (false negative). We used the *glm* function in R to fit models of the log odds of whether a non-contributing species is correlated with its corresponding metabolite (false positive or true negative), using as predictors grouped indicator values for species and metabolite identities. We separately fit another set of logistic regression models to predict whether a key contributor species is correlated (true positive or false negative), with the same predictors. Models were compared using likelihood ratio tests using the *anova* function in R.

### Simulations with varied inflow quantities

We ran 8 additional sets of simulations with the same set of 61 different initial species compositions but with varying degrees of inflow fluctuations. Specifically, the nutrient inflow quantities were sampled independently from a normal distribution, with a mean of the original inflow concentration and the standard deviation equal to a set percent of the mean. The 8 levels of deviation were 0.5%, 1%, 2%, 3%, 4%, 5%, 8%, or 10%. In the comparison of correlation results across samples, we evaluated the same set of 52 variable metabolites as for the original dataset for consistency, although given the added noise, additional metabolites met the same variance cutoff we used to define variable metabolites.

To evaluate correlation performance as a function of increasing environmental contribution, we binned the 38 analyzed inflow metabolites across the 8 datasets based on the size of the environmental contribution to variance for the metabolite in that dataset. In other words, metabolites in any dataset with an environmental contribution greater than 0 but less than 10% of the total positive variance contributions were binned into a single category, those with an environmental contribution between 10% and 20% were binned into the next category, and so on. We analyzed the 52 metabolites in the original constant-environment dataset as a separate category, and did the same for the 14 non-inflow metabolites in each of the 8 environmentally-varying datasets.

Confidence intervals for AUC values were calculated using the *pROC* package in R (78), using a bootstrap method with 500 resamplings.

### Simulations of Human Microbiome Project-based microbiota

To simulate more complex gut microbiota, we downloaded the 16S rRNA sequence variant abundance tables from the Human Microbiome Project (52), processed with deblur (79), from Qiita (80). We also downloaded ribosomal RNA sequences for all of the 818 genomes corresponding with AGORA v1.0.2 models from NCBI RefSeq and GenBank using the biomartr R package (81). We used *vsearch* version 2.8.1 (82) to map the HMP sequences to the AGORA ribosomal sequences with 97% identity, with the max_rejects parameter set to 0 in order to obtain the highest identity match for each sequence variant. We chose to model a subset of 57 samples for which at least 25% of their total read counts successfully mapped to an AGORA genome. We normalized species abundances based on the 16S rRNA copy number of the corresponding genome, and initialized 57 simulations with the starting relative abundances determined based on the AGORA-mapped relative abundances of these samples. We updated the nutrient inflow to enable growth by most models. We assessed whether the additional of each individual metabolite to the original nutrient inflow had a growth-promoting effect on any species, specifying proportions similar to the average European diet in the Virtual Metabolic Human database where possible (83). Metabolites that promoted growth in at least one species were retained in the revised nutrient inflow, and the process of testing for increased growth with the addition of any single metabolite was repeated. After two rounds of adding metabolites to the inflow, 15 models, representing 3.4% of the total normalized abundance across all samples, still displayed zero growth. We removed these from the simulations and used the final updated nutrient inflow with the 131 remaining models. All other simulation parameters were the same as for the original 10-species community simulations. When analyzing the role of interspecies correlation in this dataset, we excluded species that appear in fewer than 4 samples.

### Application of MIMOSA to simulated data and comparison with correlation analysis

We applied MIMOSA v1.0.2 (github.com/borenstein-lab/MIMOSA) (57) to the obtained set of metabolite and species abundances. To construct the community metabolic network model required by MIMOSA, we merged the 10 species-level models used in the simulations into a single stoichiometric matrix. If a reversible reaction only ever proceeded in a single direction in any simulation, we encoded it as non-reversible. To apply the KEGG-based version of MIMOSA, we converted the model metabolite IDs to KEGG IDs (56), downloaded KEGG Orthology gene annotations for the 10 modeled species from the IMG/M database (84), and ran a MIMOSA analysis using the KEGG metabolic network model encoded in *reaction_mapformula.lst* (KEGG version downloaded 2-2018).

### Code and data availability

Code for all the analyses presented in this study is available online in the form of R notebooks at https://github.com/borenstein-lab/microbiome-metabolome-evaluation. The code and media files for performing dynamic FBA co-culture simulations is available from https://borensteinlab.com/download.html. All data generated and analyzed in this study and displayed in the figures are included in Supplementary Data 1 through 4.

## Supporting information

Supplementary Results and Figures S1-11

## Author contributions

C.N. and E.B. designed the study and wrote the paper. C.N. performed the analysis. H.C.C. and C.P.M. contributed to the multi-species metabolic modeling simulations. All authors read and approved the paper.

## Acknowledgements

C.N. was supported in part by a National Science Foundation (NSF) IGERT DGE-1258485 fellowship. C.P.M. was funded by NHGRI grant T32 HG000035. This work was supported in part by NIH New Innovator Award DP2 AT007802–01 and NIH grant 1R01GM124312–01 to E.B.

