## Supplementary Results and Figures S1-11 for "Defining and Evaluating Microbial Contributions to Metabolite Variation in Microbiome-Metabolome Association Studies"

***Simulated species responses to media variation partially recapitulate experimental results***

We ran several sets of simulations with the same set of initial species compositions but we maintained the same set of 61 initial species compositions but with small amounts of stochastic noise added to the nutrient inflow, sampling inflow concentrations for each compound in each simulation from a normal distribution with a mean equal to the compound's original inflow rate and a standard deviation set to a particular fraction of the mean (Methods). We evaluated whether the differences in simulated species growth across simulations with large media fluctuations (8-10%) recapitulated the experimental observations of Faith *et al.* (47), finding agreement on several trends. First, Faith *et al.* observed that casein abundance (presumably, due to limiting amino acid or nitrogen concentrations) was strongly associated with the total community biomass. In our simulations, the total amount of amino acids is indeed positively associated with total biomass (Spearman rho=0.25, $p$=0.005). We fit linear regression models of total biomass across all simulations based on inflow metabolite levels, and found that concentrations of both amino acids and carbohydrates explain most of the variation in this quantity in our simulations (adjusted $R^2$ of 0.97), with other nutrients adding only negligible effects. Faith *et al.* further observed that while most species increase their growth rate in the presence of higher protein, *E.rectale*, *D. piger* and *M. formatexigens* are negatively associated with the amount of casein in the mouse diet. In our simulations, a model predicting growth rate based on amino acid levels explains no variation in *E. rectale* and only 8% in *M. formatexigens*, in contrast to other taxa, for which amino acids explained up to 54% of growth rate variation. Additionally, the initial growth rates for all species were positively correlated ($p < 0.1$) with an average of 2.7 different amino acid compounds (range 1 to 6), while *E. rectale* and *M. formatexigens* were each only correlated with a single one, *L*-cysteinylglycine. Lastly, our simulations also recapitulate differences in carbohydrate use, with Faith et al observing preferential expansion of *B. ovatus* and *B. thetaiotaomicron* on a high-starch diet compared to a high-sugar diet. In our simulations, the growth rate of all species was associated with the amount of available simple sugars, but only *B. ovatus*, *B. thetaiotaomicron*, and *E. rectale* were significantly correlated with the quantity of starch in the inflow (Spearman rho coefficients of 0.5, 0.53, 0.53 respectively, all $p < 10^{-9}$). These results indicate that our simulation framework successfully encapsulates some, though not all, of the nutrient limitations shaping the growth dynamics of this model community.

##### *Analysis of an alternative definition of contribution values based on flux rates*

Our contribution value metric attributes metabolite variance to each species depending on its cumulative metabolite uptake or secretion over the entire simulation, rather than its arrived-at steady-state metabolite flux at the time of "sampling". To assess the impacts of this choice, we calculated contribution values using an alternative definition based solely on steady-state fluxes. Specifically, we calculated the contribution of each species to the metabolite flux at the final time point of a simulation run for 144 hours and for 1440 hours. Under this definition, steady-state contribution values explain the variation in metabolite flux rate at the time of sampling, rather than the accumulated variation in metabolite concentrations (cumulative contribution values). We compared these alternative steady-state contributions with the original set of cumulative contribution values at both time points, finding that they are highly similar. In our original dataset of simulations run for 144 hours, the Pearson correlation between steady-state and cumulative contribution values for each metabolite was on average 0.99 (minimum of 0.75). Only 6 of the 520 analyzed species-metabolite pairs differ in contributor status between the two definitions: 4 pairs are key cumulative contributors but not steady-state contributors, and 2 pairs are the reverse. The AUC for detection of steady-state contributors is 0.710 (compared with 0.717 for cumulative contributors). These differences also naturally recede further for simulations run for longer durations: in a dataset of simulations run for 1440 hours, the average metabolite-level correlation between steady-state and cumulative contribution values was 0.999 (minimum 0.97). These results indicate that for these simulations, historical differences in species composition and metabolic activity are not a major factor in the observed discrepancy between species-metabolite correlations and true key contributors to metabolic variation.

***Analysis of an alternative definition of key taxon-metabolite pairs***

For most analyses, we defined the key taxonomic contributors for a particular metabolite as those species with the highest positive contribution values, or those that are responsible for the observed pattern of variation in a metabolite. However, an alternative goal could be to detect all microbes that substantially impact levels of a given metabolite across samples, regardless of whether their effects are ultimately reflected in the observed concentrations, To this end, we defined *key player* species as those with a contribution value, either positive or negative, greater in magnitude than 20% of the total contribution magnitude. This resulted in 91 species-metabolite key player pairs, including 65 of the previously defined 'positive' key contributor pairs but also 26 players with negative contributions, and these were distributed similarly across metabolites and species (Figure S5, panels A-B). Examining how well these key players were detected by a correlation-based analysis, we found similar performance to those reported above for key contributors (Figure S5, panels C-G), including a comparable positive predictive value (31.9%) and AUC (0.73).

##### *Effects of simulation length and* Vmax *parameter on correlation results*

We assessed the sensitivity of our correlation results to the parameters used in our simulations. Specifically, we evaluated the effect of the duration of simulations on our results. We ran additional simulations for 5,760 time points (or 1,440 hours), and calculated contribution values and correlation coefficients at 22 intermediate time points starting at 36 hours (Figure S8). Species compositions and metabolite concentrations became increasingly less variable with longer simulation time, converging towards similar steady states dominated in abundance by 5 of the 10 species (Figure S8A-B). Correspondingly, the number of key contributors decreased with increasing simulation length, from 121 contributors across all 52 analyzed metabolites at 36 hours, to 75 at 1,440 hours (Figure S8B-C). The number of significantly correlated species-metabolite pairs, however, increased from 179 to 375 over the same datasets, detecting contributors with higher sensitivity but lower specificity (Figure S8D). Ultimately, the AUC and positive predictive value both decrease slightly with increasing simulation length, with the AUC shifting from 0.67 to 0.73 and positive predictive value from 39.7% to 18.4% (Figure S8E).

This transition occurs sharply initially before reaching an inflection point and beginning to stabilize around 144 hours, the length of time chosen for our main analysis.

We also generated additional datasets with the same initial species compositions but with widely varied values for the universal *Vmax* parameter, which was set to 20 in the main set of analyses. Changing this parameter had very minimal impact on both the simulation abundance profiles and the results of correlation analysis (Figure S9). The AUC for the identification of key contributors was not associated with the value of the *Vmax* parameter, and only ranged from 0.70 to 0.72.

##### *Features distinguishing true key contributors from false positives among correlated pairs*

We constructed additional regression models to assess whether there are features that can distinguish true key contributors from false positives among all correlated species-metabolite pairs. We fit regression models to similarly assess whether species and/or metabolite identity are indicative of whether a correlated species-metabolite pair represents a true or false positive relationship. We found that species identity ($p = 0.047$), but not metabolite identity, was predictive of key contributor status among correlated pairs. This is unsurprising given that the number of key contributions from each species varied widely, while all metabolites have at least one key contributor.

##### *Additional effects of inflow fluctuations on contribution and correlation profiles*

We assessed whether the addition of external metabolite fluctuations impacted the profile of species contributing to each metabolite. For most metabolites (28 out of 52, including 12 out of 14 non-inflow metabolites), the top microbial contributor did not change across all levels of fluctuation. However, for many inflow metabolites, the large external fluctuations can result in a switch in contribution values. In these cases, activity by a microbe that contributed to variation in a constant-inflow setting instead has a mitigating impact, resulting in a negative contribution. Of the 65 key contributors to variation in inflow metabolites in the original dataset, 34 (52%) of them have a negative contribution value in at least one simulation run with external fluctuations. This observation highlights that our definition of key contributors is context-dependent, identifying the entities primarily responsible for the observed variation.

We also examined whether the detection of the 14 variable metabolites not present in the nutrient inflow was affected by random fluctuations in inflow metabolites. Variation in 8 of these metabolites was significantly positively correlated with variation in the surrounding inflow (Spearman rho, $p < 0.01$), suggesting that their synthesis fluxes were affected by changes in microbial growth or nutrient usage that resulted from environmental shifts. Correlation analysis tended to identify key contributors for these metabolites with slightly higher specificity and lower sensitivity as inflow fluctuations increased (Figure S10).

### Supplementary Figures

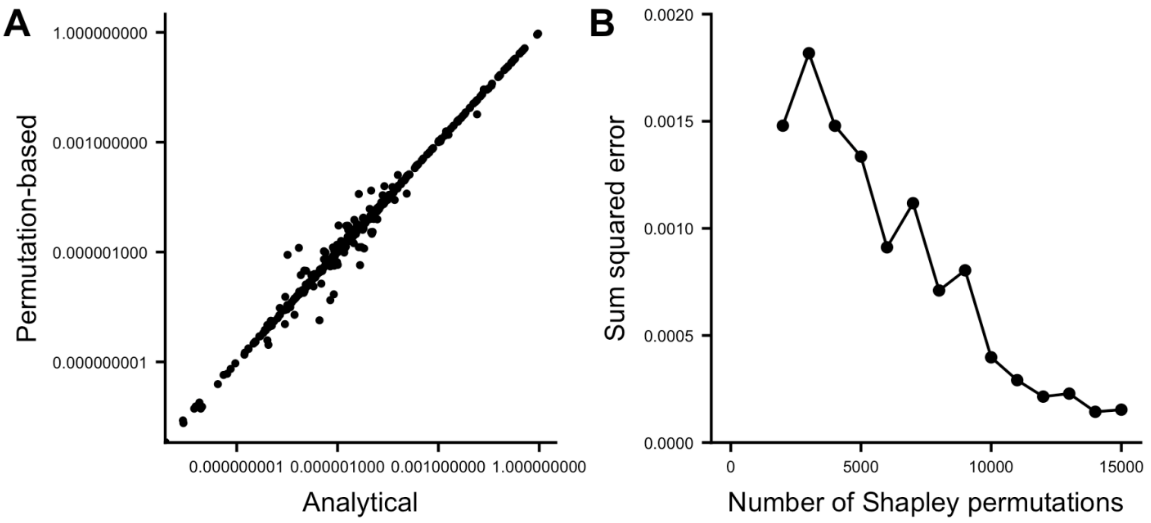


**Figure S1. Shapley values are equivalent to analytically calculated variance contributions. (A)** Plot of contribution values calculated analytically versus those obtained from a Shapley value-based permutation analysis using 15,000 species orderings (see Methods), for all 52 analyzed metabolites in our simulated dataset. Axes are on a $\log_{10}$ scale. **(B)** Plot of the total sum of squared error between Shapley values calculated using permuted species subsets and our analytically calculated variance contributions, for all metabolites. With increasing numbers of permutations and therefore increasingly precise contribution estimates, the difference between these values approaches 0.

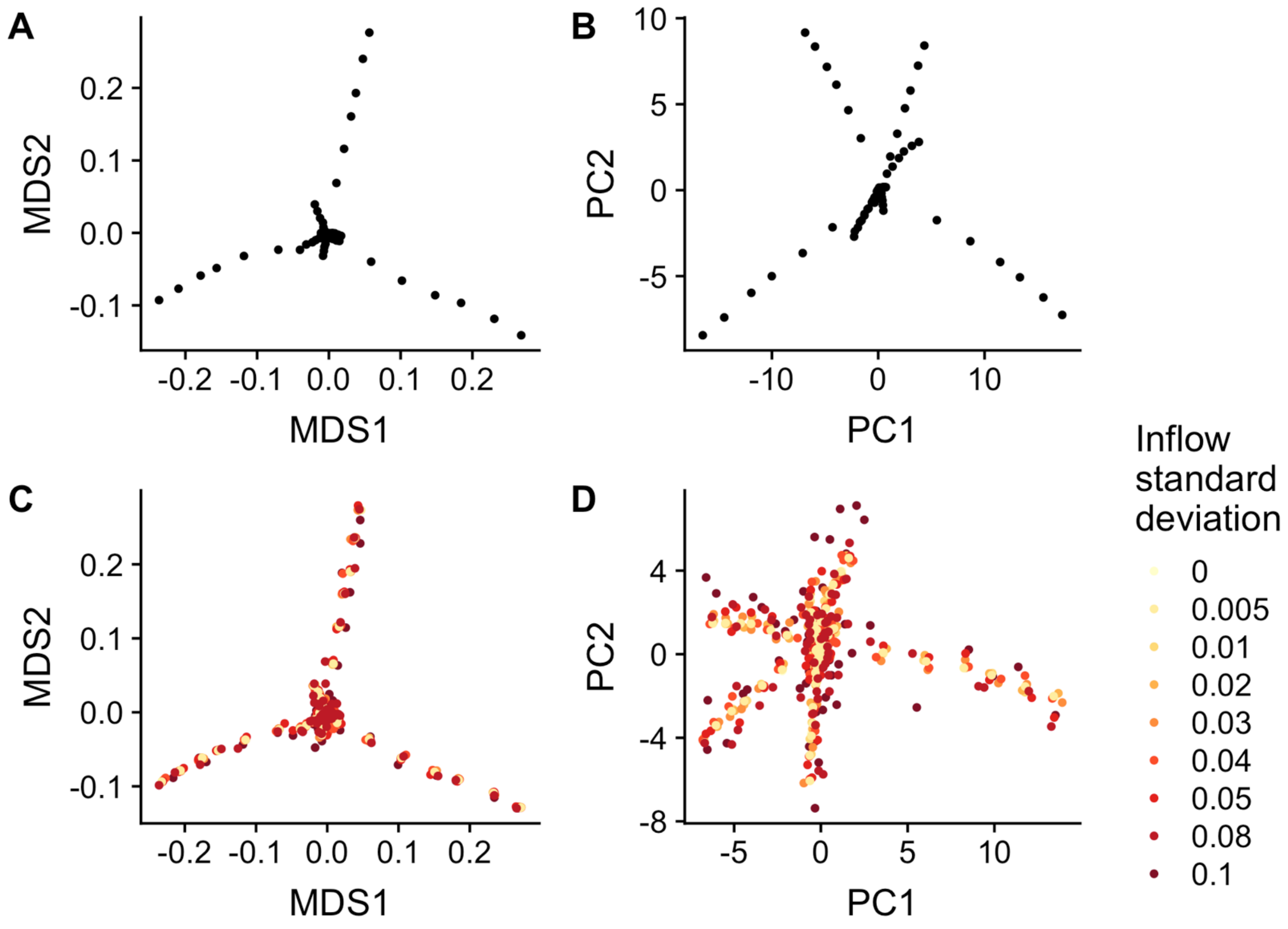


**Figure S2. Ordination plots of species and metabolite abundances in simulated datasets. (A)** Non-metric multidimensional scaling plots of species composition across the 61 original simulation runs, using Bray-Curtis dissimilarity. **(B)** Principal components analysis of metabolite concentrations across the 61 original simulation runs. **(C-D)** The same plots as **(A)** and **(B)**, but including all simulation runs with environmental fluctuations in the nutrient inflow.

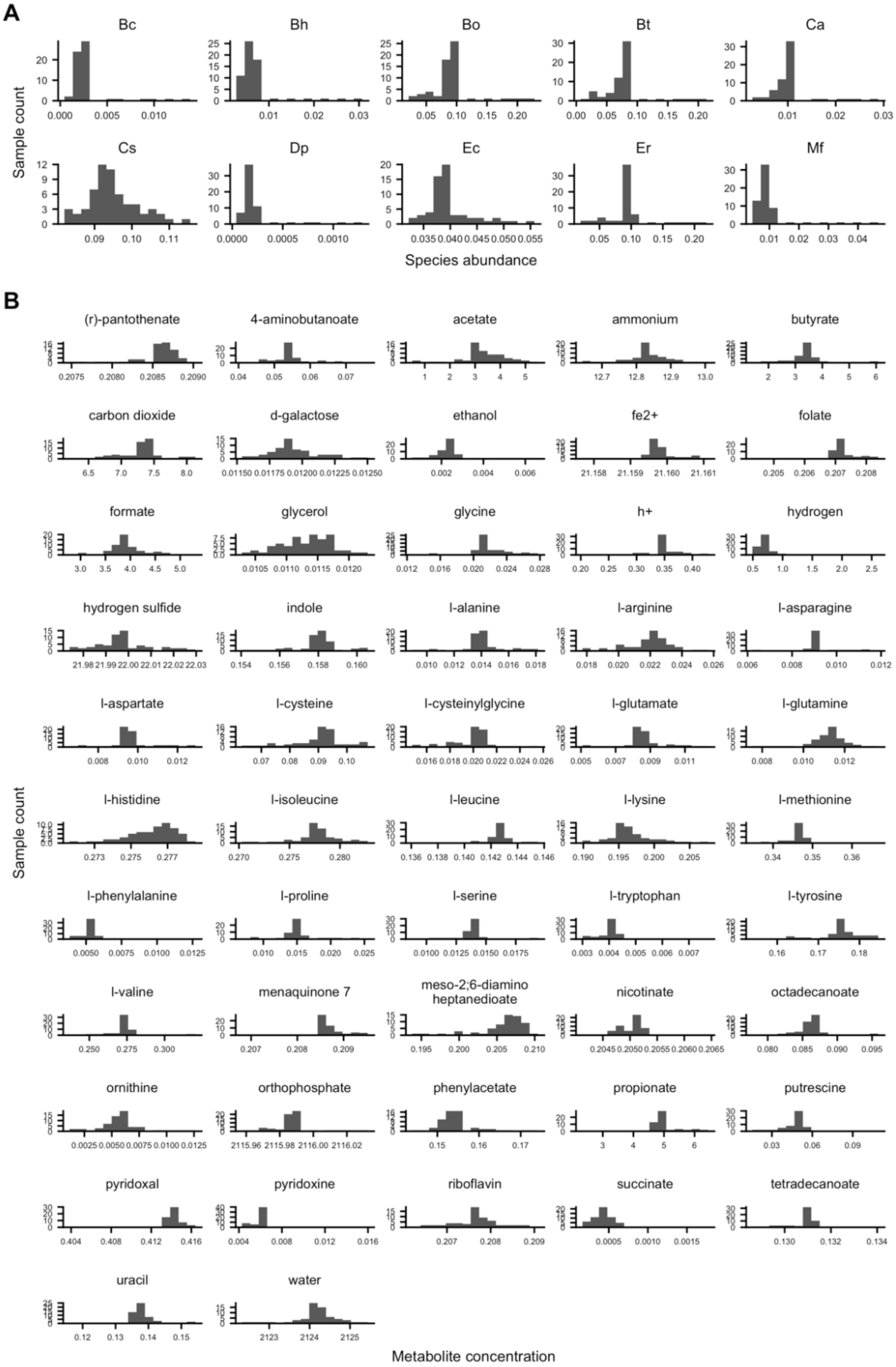


**Figure S3. Distributions of species and metabolite abundances.** Each panel shows a histogram of abundances for a single species (A) or a single variable metabolite (B) across all 61 simulation runs.

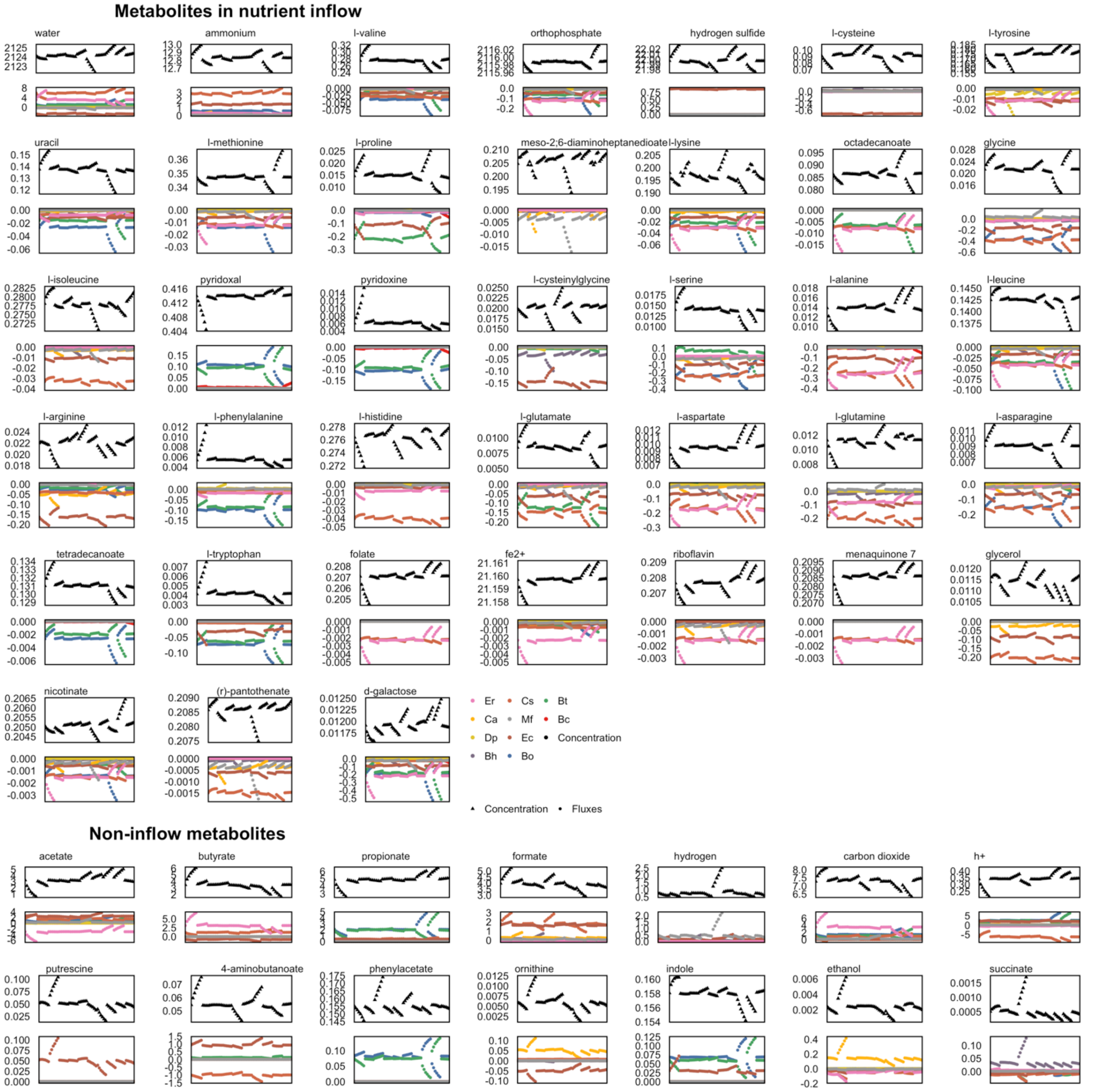


**Figure S4. Cumulative uptake and secretion fluxes for all species and all metabolites, across all 61 simulations.** For all analyzed metabolites, an upper panel shows the total cumulative secretion or uptake of that metabolite by each species across all 61 simulation runs. A lower panel shows the corresponding environmental concentration at the final time point. Each plot shows fluxes for a single metabolite, with those found in the nutrient inflow in the upper section and microbially-produced metabolites below. Metabolites are ordered by their total variance. Simulations are ordered on the x-axis in the same ordering as in Figures 1 and 2.

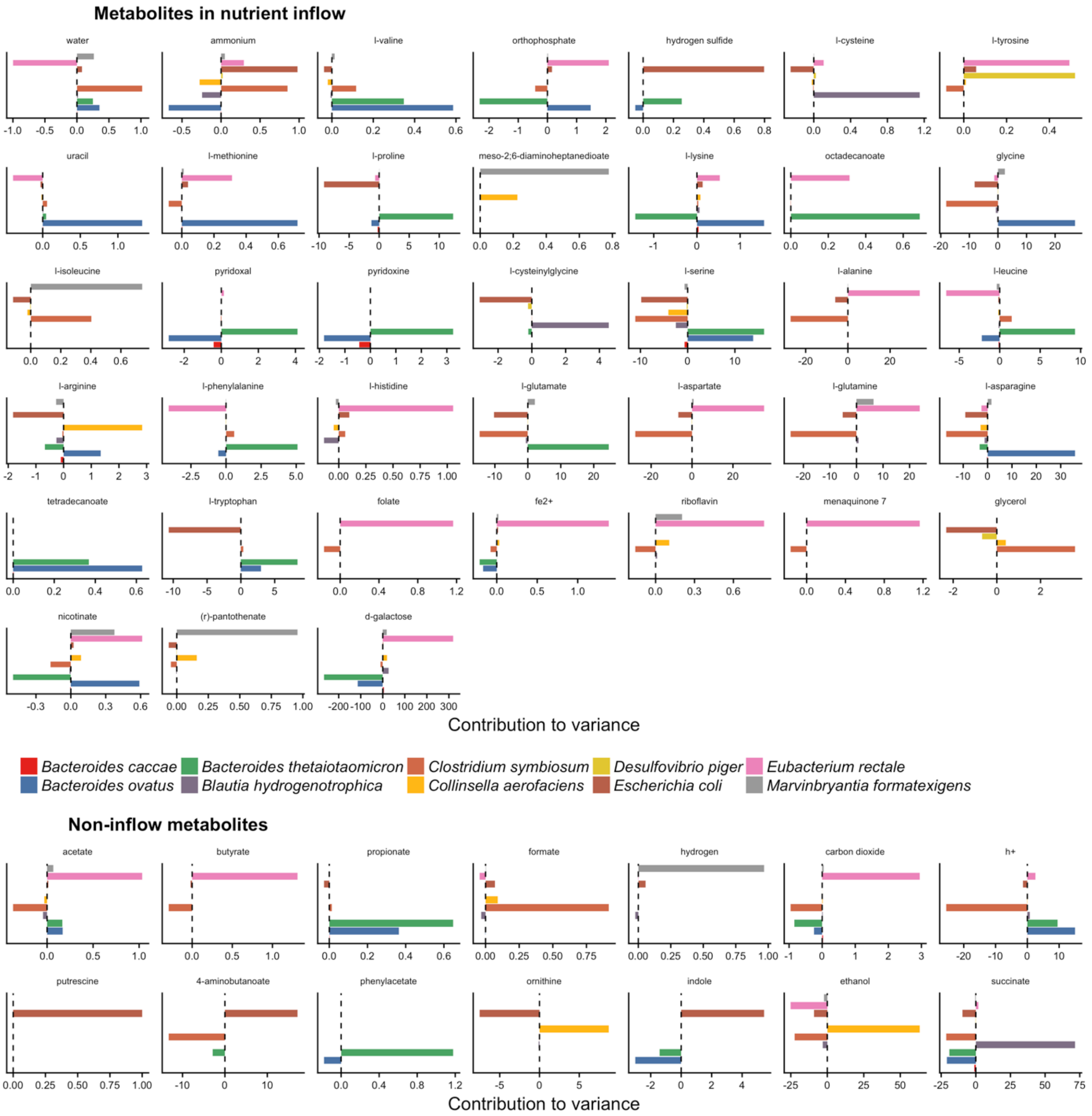


**Figure S5. Variance contribution profiles for all metabolites.** Each plot shows contribution values for a single metabolite, with those found in the nutrient inflow in the upper section and microbially-produced metabolites below. Metabolites are ordered by their total variance. The relative contribution values, $\hat{c}_i$, are plotted on the x-axis.

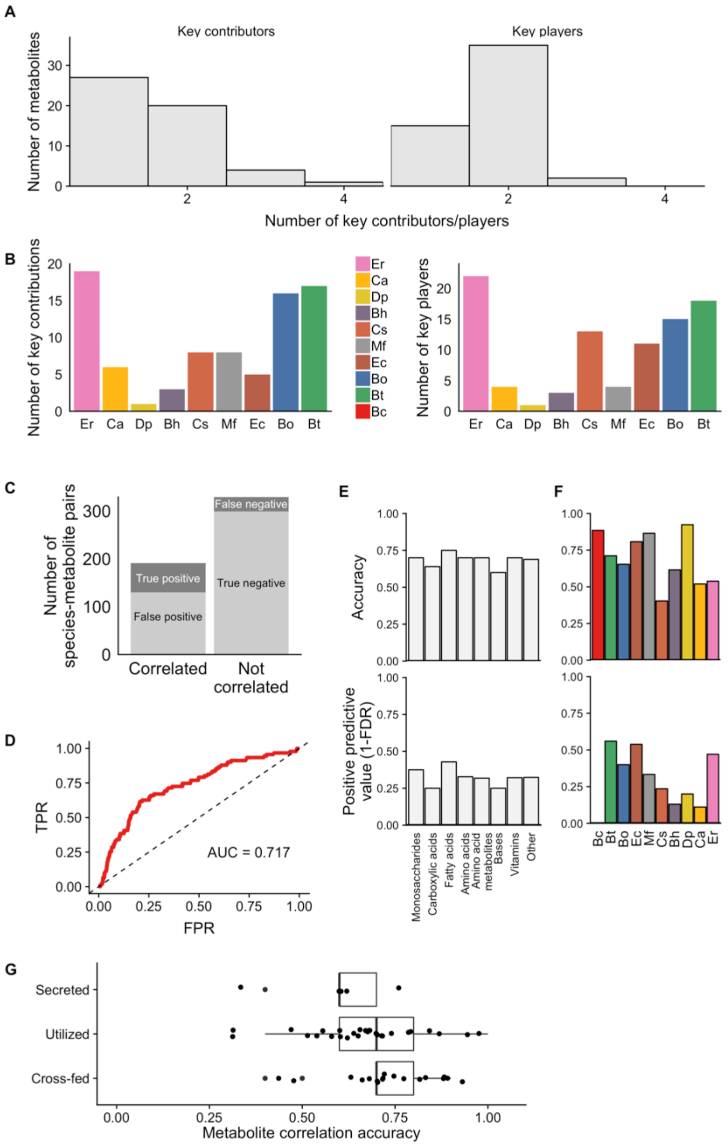


**Figure S6. Key contributors and key players driving metabolite variance have similar properties and correlation results. (A)** Histograms of the number of key contributor and key player species for all 52 analyzed metabolites. **(B)** Number of key contributor and key player relationships for each species. Full species names can be found in Figure 2. **(C-G)** Correlation results for identification of key players, as shown in Figures 3 and 4 for key contributors. **(C)** The number of species-metabolites pairs that were significantly correlated (left bar) or not correlated (right bar) and its correspondence with true species-metabolite key players. **(D)** Receiver operating characteristic (ROC) plot, showing the ability of absolute Spearman correlation values to classify key players among all species-metabolite pairs. **(E-F)** Accuracy and positive predictive value of Spearman correlation analysis for detecting true key players across metabolite classes (Panel E) and for each of the 10 species (Panel F). **(G)** As in Figure 4, correlation-based analysis detected key players equally accurately regardless of whether a metabolite is secreted, utilized, or cross-fed by the species. Each point represents the accuracy of correlation-based analysis for a single metabolite across its comparisons with all 10 species.

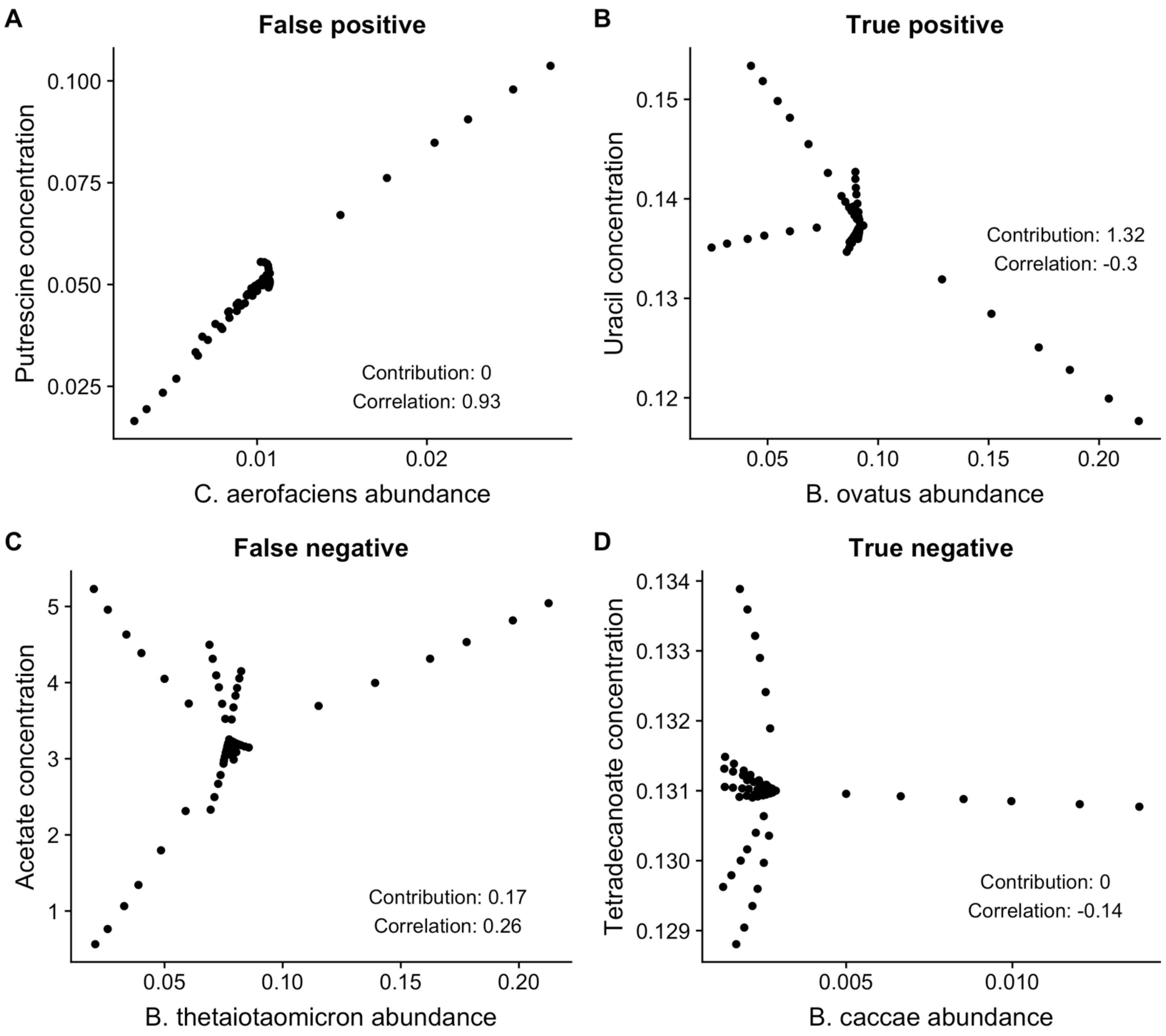


**Figure S7. Examples of species-metabolite correlation outcomes.** Each panel plots the concentration of one of the example metabolites shown in Figure 2 against the abundance of a key contributor or non-contributor species, with annotations of the corresponding correlation and contribution values.

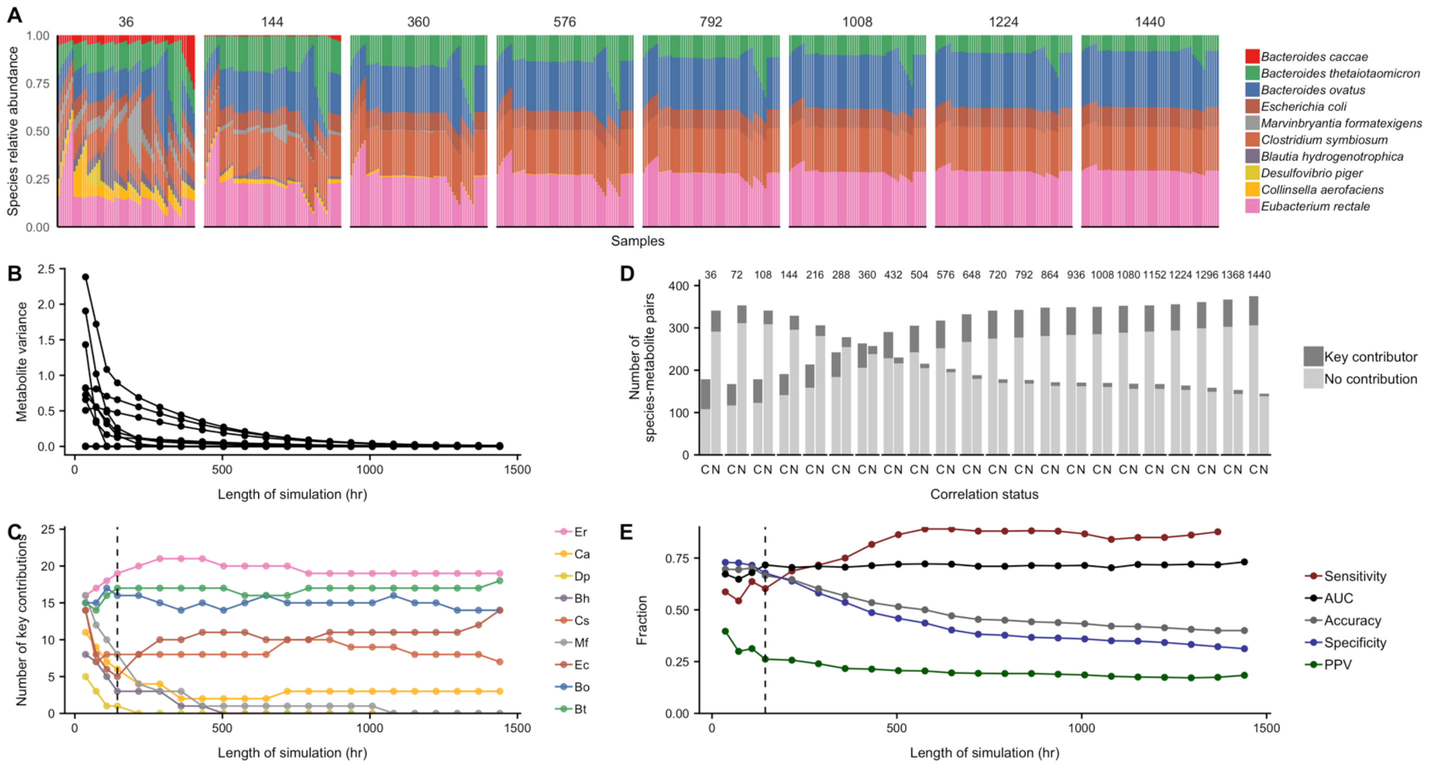


**Figure S8. Effects of simulation duration on contribution profiles and correlation efficacy. (A)** Species abundances after simulation runs of increasing length. Longer simulations increasingly converge towards a consistent profile. **(B)** Metabolite variance decreases with increasing simulation duration. Each line represents the total variance in concentration of a metabolite across the 61 simulations. **(C)** Each line represents the number of key contributions by each species across simulation datasets of increasing duration. The total number of key contributors decreases with increasing length. The length of 144 hours described in the main results is indicated with a dotted line. **(D)** Bar plots of correlation and contribution outcomes with increasing simulation duration, with the "C" labeled bar indicating the number of correlated species-metabolite pairs and the "N" indicating the number of non-correlated pairs. Datasets generated from longer simulations display more significant correlations and fewer key contributors. **(E)** Overall shifts in prediction metrics for correlation analysis with increasing simulation duration. AUC and predictive value are largely constant.

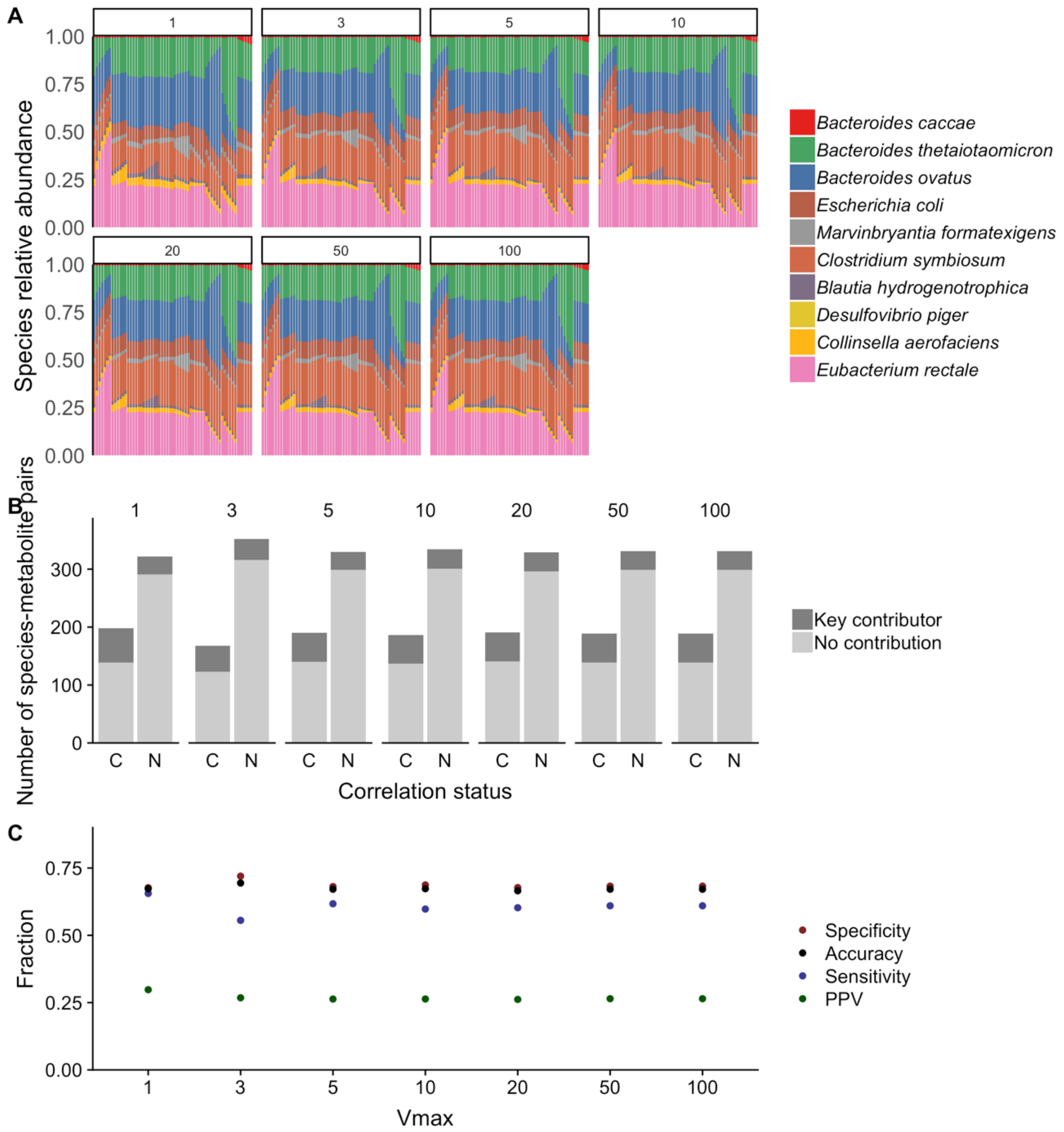


**Figure S9. Effects of *V_max* parameter on simulation and correlation results. (A)** Species compositions generated using different values of the parameter are nearly identical. **(B)** Bar plots of correlation and contribution outcomes from simulations with varying values of *V_max*, with the "C" labeled bar indicating the number of correlated species-metabolite pairs and the "N" indicating the number of non-correlated pairs. **(C)** Overall prediction metrics for correlation analysis are largely constant across simulations generated with different *V_max* values.

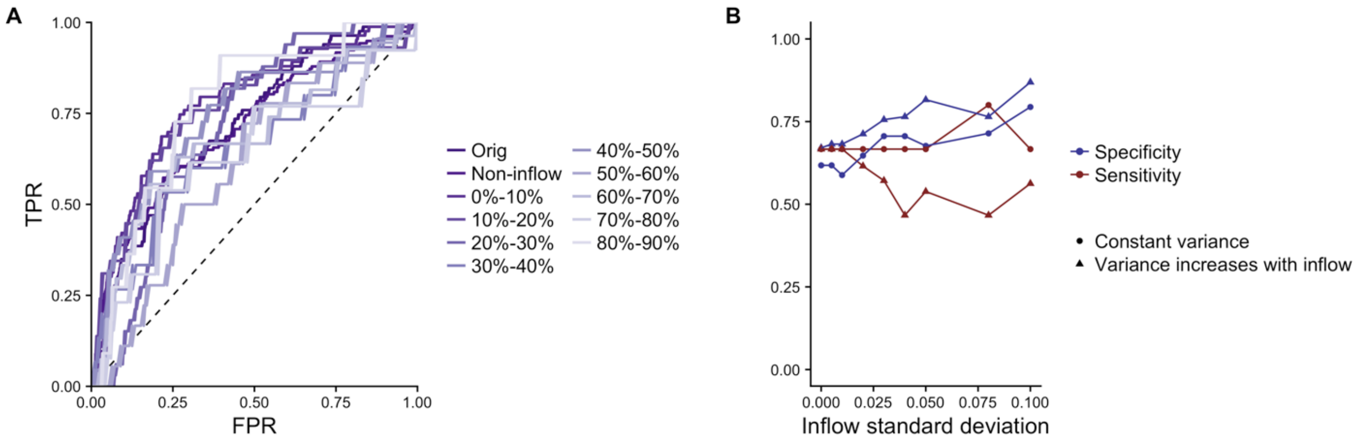


**Figure S10. (A) Environmental fluctuations do not significantly affect overall correlation performance.** ROC curves are shown for sets of metabolites with increasing environmental contribution. None of the levels of environmental contribution had a significantly different area under the curve, based on 95% confidence intervals calculated using bootstrap resampling with 500 replicates. **(B) The sensitivity and specificity of correlation analysis to detect key microbial contributors to non-inflow metabolites are affected by variation in metabolic inflow.** Each point represents the specificity, sensitivity, or positive predictive value of the 14 analyzed non-inflow metabolites in a dataset of 61 simulations. The percent standard deviation (coefficient of variation) in inflow metabolite concentrations for each set of simulations is plotted on the *x*-axis.

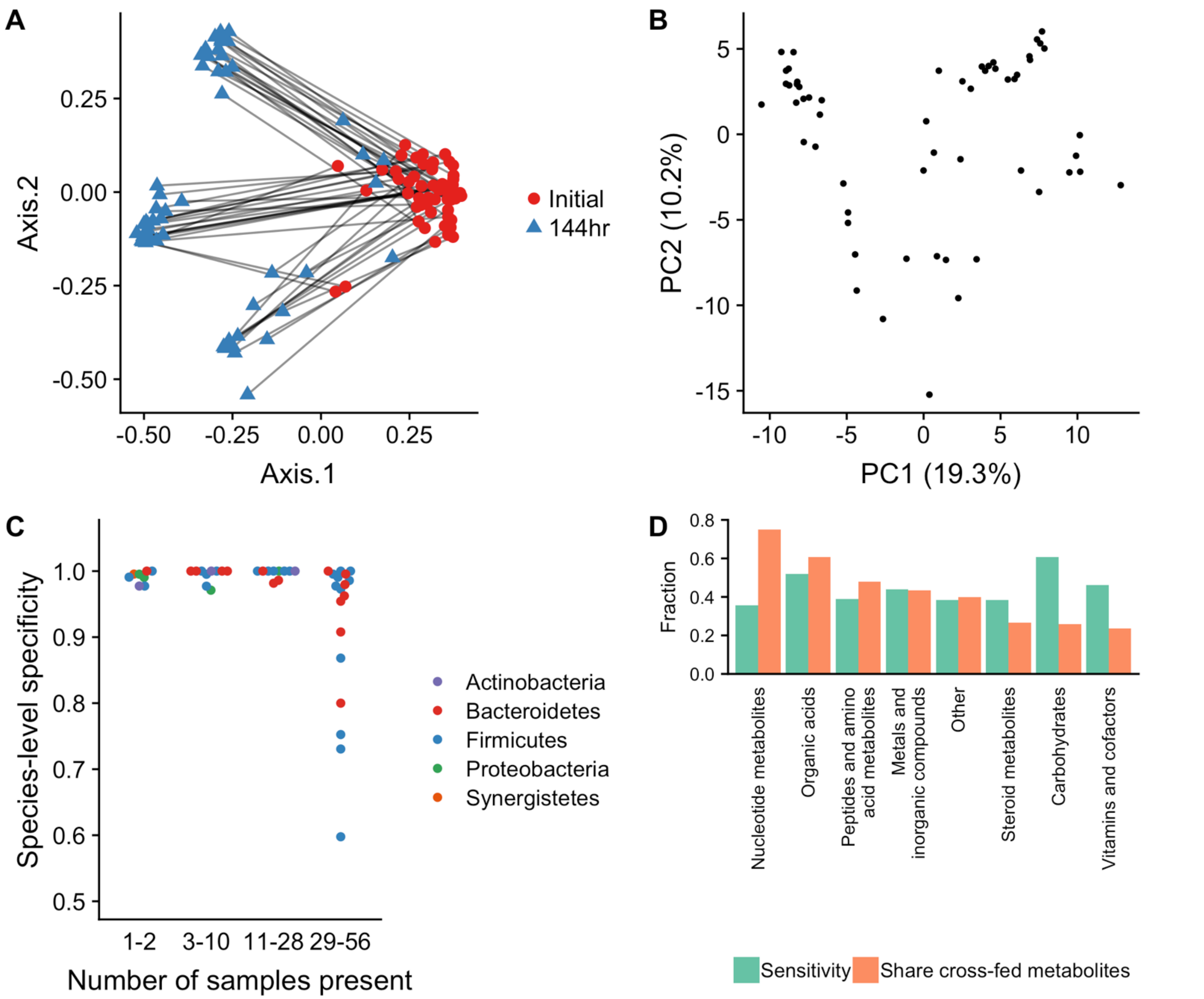


**Figure S11. (A) Progression of HMP-based simulations.** A principal coordinates analysis of the species compositions of the 57 HMP-based simulations at their initial and final time points, using the Bray-Curtis dissimilarity metric. Initial compositions tended to become dominated by a limited number of fast-growing species, leading to distinct subgroups. **(B) Metabolite variation across HMP-based simulations.** A principal component analysis of the metabolite concentration data at the final simulation time point. **(C) The specificity of species-metabolite correlation analysis is associated with species prevalence.** Each point represents a species with at least one key contribution to metabolite variation. The *x*-axis categorizes species into quartiles based on the number of samples in which they appear. Species that are present in a wider subset of the dataset have a higher rate of false positive correlations (lower specificity). **(D) The sensitivity of species-metabolite correlation analysis is related to metabolite class and cross-feeding status.** Green bars represent the overall sensitivity of identification of key contributor species-metabolite pairs within that category. Orange bars represent the share of metabolites in that category that are both synthesized and utilized by community members (cross-fed).

### Legends for Supplementary Data

**Supplementary Data 1. Simulation results and associated data, and source data for the example simulation in Figure 2.** This file contains the simulation data used for the main analysis, as well as general related parameters and metadata. **A)** The 10 genome-scale models used for all analyses. **B)** Parameter values used for the dynamic Flux Balance Analysis simulations. **C)** Nutrient inflow content, adapted from a corn-based mouse chow. **D-E)** Metabolite IDs and categorizations. **F-H)** The full simulation data at the final time step, including species abundances (F), metabolite concentrations (G), and metabolite fluxes (H). **(I-J)** Source data for Figure 2, including dynamic species abundances (I) and acetate fluxes (J) across all time steps for a single simulation.

**Supplementary Data 2. Simulation results with environment fluctuations. A-C)** The full simulation data at the final time step of each of the 8 simulations with varying levels of environmental fluctuation, including species abundances (A), metabolite concentrations (B), and metabolite fluxes (C).

**Supplementary Data 3. Simulation results for more complex microbiota. A-C)** The full simulation data at the final time step of 57 simulations based on Human Microbiome Project species composition data, including species abundances (A), metabolite concentrations (B), and metabolite fluxes (C).

**Supplementary Data 4. Reference data used for the MIMOSA analysis. A-B)** Tables used for the MIMOSA analysis, describing the mapping of species to reactions (A) and the community metabolic model of reactions and metabolites (B).
